# Neutrophilic inflammation promotes SARS-CoV-2 infectivity and augments the inflammatory responses in airway epithelial cells

**DOI:** 10.1101/2021.08.09.455472

**Authors:** BA Calvert, EJ Quiroz, Z Lorenzana, N Doan, S Kim, CN Senger, WD Wallace, MP Salomon, J Henley, AL Ryan

## Abstract

In response to viral infection, neutrophils release inflammatory mediators as part of the innate immune response, contributing to pathogen clearance through virus internalization and killing. Pre-existing co- morbidities correlating to incidence of severe COVID-19 are associated with chronic airway neutrophilia. Furthermore, examination of COVID-19 explanted lung tissue revealed a series of epithelial pathologies associated with the infiltration and activation of neutrophils, indicating neutrophil activity in response to SARS- CoV-2 infection. To determine the impact of neutrophil-epithelial interactions on the infectivity and inflammatory responses to SARS-CoV-2 infection, we developed a co-culture model of airway neutrophilia. SARS-CoV-2 infection of the airway epithelium alone does not result in a notable pro-inflammatory response from the epithelium. The addition of neutrophils induces the release of proinflammatory cytokines and stimulates a significantly augmented pro-inflammatory response subsequent SARS-CoV-2 infection. The resulting inflammatory response is polarized with differential release from the apical and basolateral side of the epithelium. Additionally, the integrity of the epithelial barrier is impaired with notable epithelial damage and infection of basal stem cells. This study reveals a key role for neutrophil-epithelial interactions in determining inflammation and infectivity in response to SARS-CoV-2 infection.

## Introduction

Novel coronavirus infectious disease, COVID-19, is caused by the severe acute respiratory distress syndrome related coronavirus 2, SARS-CoV-2 [1, 2]. While COVID-19 is associated with high hospitalization and mortality rates, a substantial proportion of the population is asymptomatic or only experiences mild symptoms. In response to viral infection neutrophils are the first and predominant immune cells recruited to the respiratory tract [3]. Neutrophils release inflammatory mediators as part of the innate immune response and contribute to pathogen clearance through virus internalization and killing [4]. While the protective versus pathological role of neutrophils in the airways during viral response is poorly understood, it has been shown that the number of neutrophils in the lower respiratory tract correlates to COVID-19 disease severity [5–7]. Infiltration of neutrophils is also characteristic of other lung diseases associated with chronic infection and inflammation, such as asthma, chronic obstructive pulmonary disease (COPD) and cystic fibrosis (CF). All these respiratory diseases have been associated with an increased risk of developing severe COVID-19 [8]. Evaluating the relationship between SARS-CoV-2 infection and pre-existing airway neutrophilia may provide critical insight into how host and viral factors contribute to disease severity.

Neutrophils have an inherent capacity to recognize infectious agents, in addition to acting as sites of infection and, in both cases, result in an acute inflammatory response [9]. Understanding the precise nature of the inflammatory response and the pathophysiological consequences, could identify pathways for therapeutic intervention based on early detection of a prognostic signature for COVID-19 outcomes. An uncontrolled, hyper-inflammatory response, known as a “cytokine storm” can result from a massive influx of innate leukocytes, inclusive of neutrophils and monocytes [10], and has been heavily implicated in patients with severe COVID-19 [11, 12]. Cytokine storm and presence of pro-inflammatory mediators can be a predictor of disease severity and often leads to acute respiratory distress syndrome (ARDS), and eventually respiratory failure [13]. Retrospective studies have also demonstrated that elevated levels of interleukin-6 (IL- 6) are a strong predictor of mortality over resolution [14], and tumor necrosis factor alpha (TNFα) is increased in severe compared to moderate cases [15].

Despite their importance in anti-viral immunity and response to viral pathogens, neutrophils have been somewhat overlooked for their role in the pathogenesis of SARS-CoV-2 infection [16–18]. It has been shown that the number of neutrophils in the lower respiratory tract correlates to disease severity in other viral infections, including influenza A infection [19] and, more recently, to also be a feature of COVID-19 pathology [18]. Several studies have highlighted the importance of neutrophils in the response to SARS-CoV-2 infection [17, 18, 20, 21] and clinically neutrophil-lymphocyte ratios (NLR) are becoming an important hallmark of severe COVID-19 [22]. Furthermore, the expression of angiotensin converting enzyme 2 (ACE2) on neutrophils has also been demonstrated [23–26]. These studies, however, have primarily focused on the recruitment of neutrophils post-infection and the production of neutrophil extracellular traps and lack insights into the infection of airways with pre-existing neutrophilia and other neutrophil functional responses such as inflammatory cytokine production and viral internalization.

In this study, the relationship between SARS-CoV-2 infection and pre-existing airway neutrophilia in differentiated airway epithelium was evaluated through the adaption of a co-culture infection model previously used to study viral infections *in vitro* [27]. Primary neutrophils were isolated from peripheral blood and co- cultured with differentiated primary tracheo-bronchial airway epithelium prior to infection with live SARS-CoV- 2 virus for 4 hours to characterize the earliest stages of infection. Changes in the inflammatory profile and epithelial response were comprehensively evaluated to determine the impact of pre-existing neutrophilia on SARS-CoV-2 infection of the airway epithelium.

## Materials and Methods

### Isolation of neutrophils from peripheral blood

Neutrophils were isolated from fresh human peripheral blood with patient consent and approval of the Institutional Review Board (IRB) of the University of Southern California (USC), protocol #HS-20-00546. CD15-expressing neutrophils were isolated using the EasySep^TM^ direct neutrophil isolation kit (Stem Cell Technologies, Seattle, WA) within 1 hour of the blood draw as per the manufacturer’s instructions. Briefly, 5 ml of peripheral blood was collected into 10 ml EDTA vacutainers (Becton Dickinson, Franklin Lakes, NJ). From this, 3 ml was diluted 1:1 with PBS (Thermo Fisher Scientific, Waltham, MA) and kept on ice for purity analysis by flow cytometry. The remaining 2 ml was transferred to a 5 ml polystyrene round bottomed tube (Genesee Scientific, San Diego, CA) and gently combined with 100 μl of isolation cocktail and 100 μl of RapidSpheres^TM^ (Stem Cell Technologies). After incubation at room temperature for 5 mins, 1.8 ml of 1 mM EDTA was added, gently mixed, and placed into the EasySep^TM^ Magnet (Stem Cell Technologies) for 5 mins. The enriched cell suspension was placed into the EasySep^TM^ Magnet for an additional 5 mins and decanted into a fresh tube. Approximately 4.25 x 10^6^ cells were isolated from 5 ml of peripheral blood.

### Flow activated cell sorting (FACS)

To validate the purity of neutrophils isolated from peripheral blood; 1x10^7^ CD15^+^ freshly isolated human neutrophils were resuspended in 100 ul FACS buffer (PBS, 0.5mM EDTA, 1% FBS, 0.1% BSA) and fresh whole human blood diluted 1:5 in FACS buffer and supplemented with 5 ul of human TruStain Fc receptor blocker (Biolegend, San Diego, CA) for 5 mins on ice. Cells were then incubated with anti-human CD15 PE (Biolegend) for 1 hour prior to FACS analysis. Cells were analyzed on the SORP FACS Symphony cell sorter (BD Biosciences) in the Flow Cytometry Facility at USC using FACS Diva software and all analyses was carried out in Flow Jo V10.8.0 (BD Biosciences).

### Air-liquid interface (ALI) differentiation of airway epithelium

Primary human airway basal epithelial cells (HBECs) were isolated from explant human lung tissue as previously described [28] and with approval of IRB at USC (protocol #HS-18-00273). For this study, HBEC donors were randomly paired with blood neutrophil donors (detailed in **supplemental table S2&3**). HBECs were expanded for 1 to 4 passages in airway epithelial cell growth media (AEGM, Promocell, Heidelberg, DE) and transitioned to Pneumacult Ex+ (Stem Cell Technologies) for 1 passage, prior to growth on Transwells. Cells were routinely passaged at 80% confluence using Accutase^TM^ (Stem Cell Technologies) and seeded at 5 x 10^4^ cells per 6.5 mm polyethylene (PET) insert with 0.4 µm pores (Corning, Corning, NY). Media was changed every 24-48 hours and transepithelial electrical resistance (TEER) was monitored every 24-48 hours using an EVOM3 epithelial volt-ohm meter (World Precision Instruments, Sarasota, FL). At resistances ≥ 450Ω ^cm^2^, cells were air lifted by removing the apical media and washing the apical surface with phosphate buffered saine (PBS, Sigma-Aldrich, St Louis, MO). The basolateral media was replaced with Pneumacult ALI media (Stem Cell Technologies) and changed every 2 to 3 days for up to 40 days.

### SARS-CoV-2 culture

Vero E6 cells overexpressing ACE2 (VeroE6-hACE2) were obtained from Dr. Jae Jung and maintained in DMEM high glucose (Thermo Fisher Scientific, Waltham, MA), supplemented with 10% FBS (Thermo Fisher Scientific, Waltham, MA), 2.5 µg/ml puromycin (Thermo Fisher Scientific, Waltham, MA) at 37^°^C, 5% CO2 in a humidified atmosphere in the Hastings Foundation and The Wright Foundation Laboratories BSL3 facility at USC. SARS-CoV-2 virus (BEI resources, Manassas, VA) was cultured and passaged 4 times in VeroE6- hACE2 cells and harvested every 48 hours post-inoculation. Plaque forming units (PFU) were determined using a plaque assay by infecting a monolayer of VeroE6-hACE2 cells with serial dilutions of virus stocks and layering semi-solid agar. Plaques were counted at day 3 post infection to determine PFU. Virus stocks were stored at -80^°^C.

### SARS-CoV-2 infection

Differentiated airway epithelium at ALI was cultured with addition of 50 µl of PBS to the apical surface and incubated at 37°C, 5% CO2 in a humidified atmosphere. After 10 minutes PBS was removed to eliminate the mucus build-up on the apical surface. The basolateral culture media was removed and replaced with 400 µl of assay media (Bronchial Epithelial Growth Media (BEGM), Lonza, Walkersville, MA), without the addition of bovine pituitary extract, hydrocortisone & GA-1000, for 1 hour prior to the addition of neutrophils. Freshly isolated neutrophils were diluted to 5 x10^6^ cells/ml in Hank’s Balanced Salt Solution (with Mg2^+^ and Ca2^+^) (Thermo Fisher Scientific, Waltham, MA) and 20 µl of this suspension was seeded onto the apical surface of the ALI cultures. Monocultures of airway epithelium and neutrophils were used as controls. The neutrophil-epithelial co-cultures were incubated for 1 hour during which they were transferred to the BSL3 facility for infection. Co-cultures were infected with 1x10^4^ PFU of SARS-CoV-2 in 100 µl of OptiMEM (Thermo Fisher Scientific, Waltham, MA) added to the apical surface. Infected cell cultures were incubated for 4 hours at 37°C, 5% CO2 in a humidified atmosphere. After infection, the apical and basolateral supernatants were collected, and SARS-CoV-2 was inactivated with 1% Triton-X (Sigma-Aldrich, Burlington, MA) in PBS for 1 hour. Culture supernatants were stored at -20°C until required.

### Validation of virus inactivation

SARS-CoV-2 virus was inactivated by addition of 10% Triton-X to supernatants to generate a final concentration of Triton-X of 1% and incubating at room temperature for 1 hour. PFU was quantified using a plaque forming assay with ACE2 over-expressing Vero E6 cells (VeroE6-hACE2). Serial dilutions of SARS- CoV-2 virus were performed from a stock concentration of 1x10^5^ PFU/ml and inactivated with 1% Triton-X at room temperature for 1 hour and used to infect Vero E6 cells for a total of 4 days. Cells were monitored routinely for cytopathic effects using the Revolve microscope (Echo Laboratories, San Diego, CA).

### RNA isolation and qRT-PCR

RNA was collected in 100 µl of Trizol (Thermo Fisher Scientific, Waltham, MA) per insert and incubated for 15 mins at room temperature. Cell isolates were gently mixed by pipetting up and down. An additional 900 µl of Trizol was added and cell isolates were collected and stored at -80°C until required. Cellular RNA was isolated by either phenol/chloroform extraction or using the Direct-zol RNA Microprep kit (Zymo Research, Irvine, CA). RT-qPCR was performed in 384 well plates on an Applied Biosystems 7900HT Fast Real-Time PCR system using the QuantiTect Virus Kit (Qiagen, Redwood City, CA) and SARS-CoV-CDC RUO primers and probes (Integrated DNA Technologies (IDT), Coralville, IA). Briefly, each 5 µl reaction contained 1 µl 5x QuantiTect Virus Master Mix, 500 nM forward primer, 500 nM Reverse Primer, 125 nM Probe, 10 ng DNA, 0.05 µl QuantiTect Virus RT Mix, and DNAse/RNAse-free water up to a final volume of 5 µl. Calibration curves for RNAseP primers/probe was performed with 10-fold dilutions of RNA from uninfected Calu3 cells (ATCC, Manassas, VA) from 100 ng to 0.01 ng per reaction. Calibration curves for N1 primers were performed on 5 ng of RNA from uninfected Calu3 cells per reaction spiked with 10-fold dilutions from 50 ng to 0.005 ng of RNA from Calu3 cells collected 48 hours post infection. Relative gene expression was calculated using the Pfaffl method [29].

### Immunohisto-/cyto-chemistry

Primary human lung tissue from post-mortem or surgical resection donors (detailed in **supplemental table S1**) was fixed in 10% neutral buffered Formalin (Thermo Fisher Scientific, Waltham, MA). The tissue was then dehydrated in 70% ethanol (Thermo Fisher Scientific, Waltham, MA) prior to embedding in paraffin blocks for sectioning. Tissue sections were mounted on positively charged slides (VWR, Visalia, CA) and tissue was rehydrated through sequentially decreasing concentrations of ethanol (100% - 70%) and finally water. Slides were stained sequentially with Hematoxylin and then Eosin and imaged on the Olympus microscope IX83 (Olympus, Waltham, MA). Alternatively, tissue slides were incubated overnight at 60°C in Tris-based antigen unmasking solution (Vector Laboratories, Burlingame, CA) before permeabilization in 3% BSA, 0.3% Triton- X 100 in PBS for 1 hour and blocking in 5% normal donkey serum (Jackson ImmumoResearch, West Grove, PA) for 1 hour at room temperature. *In vitro* co-cultures were fixed in 4% PFA (Thermo Fisher Scientific, Waltham, MA) for 1 hour at room temperature and stored in PBS at 4°C to be used for immunohisto/cytochemistry. Co-cultures were then permeabilized and blocked in 3% BSA, 0.3% Triton-X 100 in PBS for 1 hour and blocking in 5% normal donkey serum (Jackson ImmumoResearch, #017-000-121) for 1 hour at room temperature. Tissue sections and *in vitro* cultures were subsequently stained with the antibodies or RNAScope probes listed in **supplemental table S4**. Slides were mounted in Fluoromount-G (Thermo Fisher Scientific, Waltham, MA) and imaged on a DMi8 fluorescent microscope (Leica, Buffalo Grove, IL) or a Zeiss LSM 800 confocal microscope (Zeiss, Dublin, CA).

### Transepithelial Electrical Resistance

Pre-warmed assay media (200 μl) was added to the apical surface of the cultures and TEER was measured using an EVOM-3 meter (World Precision Instruments).

### Meso Scale Discovery cytokine assay

50 μl of apical and 50 μl basolateral cell culture supernatants were analyzed for cytokines using the Meso Scale Discovery (MSD) V-plex Viral Panel 1 Human Kit (Meso Scale Diagnostics, Rockville, MA) as per the manufacturer’s instructions. Briefly, 1:5 dilutions of cell supernatant samples were diluted in PBS containing 1% Triton-X. Samples were added to the MSD plate along with a 7-point 4-fold serial dilution (concentrations related to certificate of analysis for each individual standard) of protein standards diluted in PBS with 1% Triton-X. The MSD plate was sealed, and samples incubated at room temperature for 2 hours on a plate shaker (ThermoFisher Scientific, Waltham, MA) at 700RPM. The plate was washed 3x in wash buffer and 25 μl of secondary antibody was added to each well. Plates were sealed and incubated at room temperature on a plate shaker at 700RPM for a further 2 hours in the dark. Plates were washed 3x with wash buffer and 50 μl of 2x read buffer (MSD R92TC) was added to each well. The plates were read on the MESO Sector S 600 (Meso Scale Diagnostics) and concentrations determined against the standard curves.

### Meso Scale Discovery SARS-CoV-2 Spike protein assay

25 μl of apical and 25 μl basolateral cell culture supernatants were analyzed for cytokines using the MSD S- plex SARS-CoV-2 Spike Kit as per the manufacturer’s instructions. Briefly, plates were washed 3x in wash buffer (PBS 0.05% Tween-20) and coated with 50 μl of coating solution (1:40 dilution of Biotin SARS-CoV-2 spike antibody; 1:200 dilution of S-PLEX Coating reagent C1 in Diluent 100) and incubated at room temperature on a plate shaker at 700RPM for 1 hour. Plates were then washed 3x in wash buffer and blocked in 25 μl blocking solution (1:100 dilution of Blocker s1 in Diluent 61) per well. Samples were added to the MSD plate along with a 7-point 4-fold serial dilution (concentrations related to certificate of analysis for each individual standard) of protein standards diluted in PBS with 1% Triton-X. Plates were incubated at room temperature on a plate shaker at 700RPM for 1.5 hours. Plates were washed 3x in wash buffer and 50 μl per well of TURBO-BOOST antibody (1:200 dilution of TURBO-BOOST SARS-CoV-2 Spike antibody in Diluent 59) was added to each well and plates were incubated at room temperature on a plate shaker at 700RPM for 1 hour. Plates were washed 3x in wash buffer and 50 μl per well of Enhance Solution (1:4 dilution of S-plex Enhance E1 1:4 dilution of S-plex Enhance E2 and 1:200 dilution of S-plex Enhance E3 in molecular biology grade water) was added. Plates were incubated at room temperature on a plate shaker at 700RPM for 30 mins. Plates were washed 3x in wash buffer and 50 μl of Detection solution (1:4 dilution of S-plex Detect D1 and 1:200 dilution of S-plex detect D2 in molecular biology grade water) was added to each well. Plates were incubated at 27°C on a plate shaker at 700RPM for 1 hour. Plates were washed 3x in wash buffer and 150 μl on MSD GOLD Read Buffer B was added to each well. Plates were read immediately on a MSD 1300 MESO QuickPlex SQ 120 plate reader (Meso Scale Diagnostics) and concentrations determined against the standard curve.

### Viral Internalization Assay

CD15+ neutrophils were seeded at 20,000 cells per well in in HBSS with or without 15 μM Cytochalasin D (Sigma Aldrich, Burlington) black walled 96 well plates (Thermo Fisher Scientific, Waltham, MA) for 1 hour to allow for attachment. Cells were then infected with SARS-CoV-2 at 2 MOI (80 μl at 5x10^5 PFU/ml) for 4 hours. Cells were then washed 2 x with PBS and fixed in 4% PFA. Cells were stained for SARS-CoV-2 RNA via RNAScope and DAPI as per the manufacturer’s instructions. Whole wells were supplemented with 50 μl of PBS post staining and well were scanned on the DMi8 fluorescent microscope (Leica, Buffalo Grove, IL)). Total cell number was determined by total frequency of DAPI particles and infected cells determined by SARS-CoV-2 particle signal in proximity to DAPI. Images were analyzed with ImageJ software 1.52n (National Institute of Health, Bethesda, MA).

### Data Analysis and Statistics

All data are presented as mean ± S.E.M. Statistical analysis is dependent upon the data set and is specifically indicated in each figure. For comparisons of 2 groups. a two-tailed unpaired Student’s T-test was used. For more than 2 groups, an analysis of variance (ANOVA) was used with a post hoc Tukey test. Significance is determined to be P<0.05. All data represents a minimum of three independent biological replicates (N=3), each with 3 experimental replicates (n=3). Data was presented and analyzed using Graph Pad prism v8.4.3 (GraphPad, San Diego, CA).

## Results

### In vitro models of neutrophilic airways have significant, polarized inflammatory responses to SARS- CoV-2 infection

Given the prevalence of neutrophilia in the airways of patients with chronic airway disease [30] and its association with other SARS-CoV-2 co-morbidities, such as diabetes mellitus [31] and hypertension [32, 33], the impact of chronic neutrophilic airway inflammation in the initial stages of SARS-CoV-2 infection was evaluated. We adapted a neutrophilic airway *in vitro* model, previously described by Deng and colleagues [27], co-culturing CD15^+^ peripheral blood polymorphonuclear leukocytes (PMNs) with primary HBECs differentiated at the ALI and infected these cultures with live SARS-CoV-2 virus for 4 hours, shown in the schematic in **figure 1a**. This 4-hour time point allows for profiling of the initial stages of infection and acute phase cellular viral response, i.e., neutrophil degranulation. The short time frame for analysis was chosen to eliminate significant viral replication and thus anticipate any detectible intracellular viral load is as a result of initial infection [34], and to allow for optimal investigation into neutrophil function without loss of viability interfering with the assays due to the relatively short half-life of neutrophils. Prior to infection we confirmed the expression of ACE2 and Transmembrane Serine Protease 2 (TMPRSS2) in our *in vitro* airway epithelium models (**supplementary figure S1)**. While ACE2 RNA was relatively low in expression across basal, secretory and multiciliated cells (**supplementary figure S1a-c**) at the protein level a predominant colocalization was detected with multiciliated cells in the airways (**supplementary figure S1a, d-f)).** This data supported by similar analysis of human lung tissues (**supplementary information and supplementary figure S2**) where we observed a similarly low level of expression in RNA in basal, secretory and multiciliated cells (**supplementary figure S2a-b**) while protein, detected by IF, was associated with multiciliated cells and cells in submucosal glands (**supplementary figure S2c-f**). Confirmation of ACE2 expression at the RNA and protein level in human lung tissues and our *in vitro* model supports currently published data evaluating ACE2 in human lung tissue [35–37].

**Figure 1:**
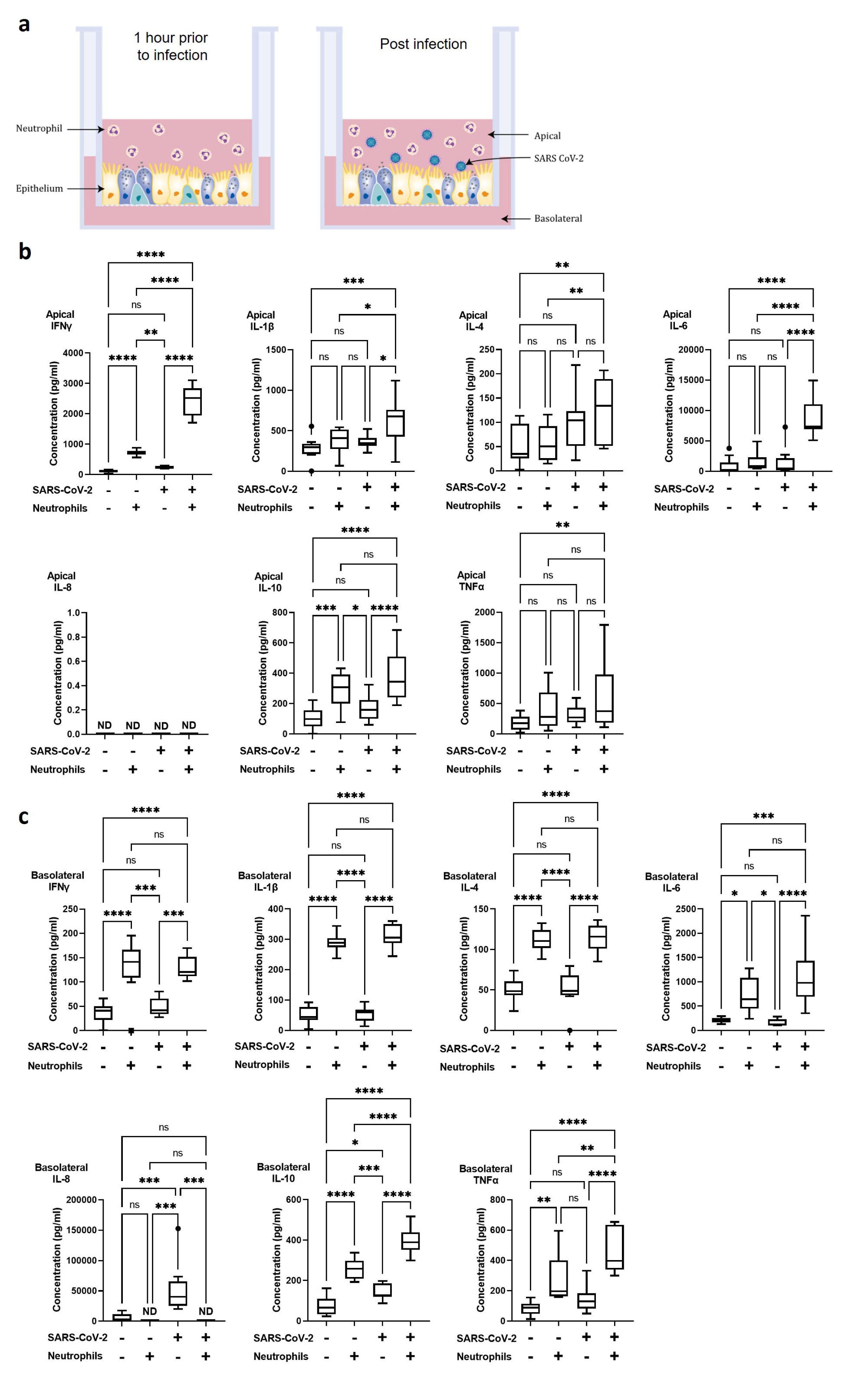
Polarized inflammatory response of neutrophils in co-culture with human airway epithelium, infected with SARS-CoV-2. a) Schematic of the *in vitro* model of neutrophilic airways denoting neutrophils in co-culture with differentiated airway epithelial cells and infected with live SARS-CoV-2 virus. Inflammatory profiles of apical (b) and basolateral (c) supernatants collected 4 hours post infection in the neutrophilic airway model. Data is expressed as Tukey method box & whiskers plots. Significance is determined by analysis of variance (ANOVA) followed by Tukey’s post hoc analysis. *p<0.05, **p<0.01, ***p<0.001, ****p<0.0001 from n=3 experimental repeats from N=3 biological donors.

In our model system the apical side of the epithelium comprises predominantly multiciliated and secretory cells directly exposed to neutrophils and the virus, the basolateral side predominantly comprises of basal cells. To understand the immediate inflammatory response of the airway epithelium to SARS-CoV-2 infection we evaluated both the apical and basolateral cell culture supernatants using the meso scale discovery (MSD) cytokine assay. All experiments were carried out using three independent HBEC donors and three independent neutrophil donors ensuring significant biological variability in our model system. As shown in **figure 1b&c** a differential inflammatory profile exists between the apical and basolateral compartments. Focusing first on the apical cytokine and chemokine release, in the absence of neutrophils, there were, surprisingly. no significant changes in cytokine release from the airway epithelial cells upon SARS-CoV-2 infection (**figure 1b**). The addition of neutrophils to the model, creating a neutrophil-epithelial co-culture in the absence of any infection, resulted in a significant secretion of interferon gamma (IFNγ, 634±1.6%, p<0.01) and IL-10 (273±11.6%, p<0.01) at the apical surface with notable, but not statistically significant, increases in tumor necrosis factor alpha (TNFα) (**figure 1b**). Like the apical release, in the airway only cultures only basolateral release of interleukin-8 (IL-8), which increased from 6180±1751 to 52996±17121 pg/ml, p<0.001, and basolateral release of IL-10, which increased from 74.42±15.36 to 142.4±12.82 pg/ml (p<0.05), were significantly changed in response to SARS-CoV-2 infection (**figure 1c**). As IL-8 is a major chemoattractant for neutrophils this suggests that the basolateral surface responds to viral infection by releasing IL-8 to recruit neutrophils to infection site [38–40]. The addition of neutrophils to the airway stimulated the release of IFNγ (321±4.1%, p< 0.05) and IL-10 (341±8.5%, p<0.01) and additionally significantly increased the release of IL1- β (557±4%, p<0.0001), IL-4 (220±3%, p<0.0001), IL-6 (761.9 ± 120.7, p<0.05) and TNFα (274 ±57.8, p<0.01) from the basolateral surface (**figure 1c**). Interestingly, the presence of neutrophils did not stimulate significant changes in IL-8 secretion from the basolateral surface supporting the role for IL-8 in the recruitment phase of airway neutrophilia, already established in our neutrophilic airway model (**figure 1c**) [41, 42]. This data demonstrates that a pro-inflammatory niche is driven primarily by the neutrophils, likely though degranulation. Based on this information we added neutrophils to our airway epithelium to create a pro-inflammatory niche recreating aspects of chronic airway inflammation in the human lung in an *in vitro* model.

Infection of the neutrophilic airway models with live SARS-CoV-2 virus was compared directly to both the infection in the absence of neutrophils and the neutrophilic airway in the absence of infection. Changes in inflammatory cytokine release from both the apical and basolateral surfaces was significantly augmented compared to both the infected epithelial monocultures and the non-infected co-cultures, demonstrating an exacerbation of pro-inflammatory cytokine release in the infected co-cultures (**figure 1b&c).** Compared to the infected epithelial monocultures, infection of the co-culture model resulted in a significant increase in the apical secretion of IFNγ, IL1-β, IL-6 and IL10 (1030±5%, p<0.0001; 169±6%, p<0.05; 580±8% p<0.0001 and 231±3%, p<0.001, respectively) (**figure 1b**) and in the basolateral secretion of IFNγ, IL1-β, IL-4, IL-6, IL10 and TNFα (261±5%, p<0.05; 572±6% p<0.0001; 203±5% p<0.001; 593±8% p<0.0001; 279±8% p<0.001 and 316±2%, p<0.001, respectively) (**figure 1c**). Compared to the uninfected neutrophil-epithelial co-cultures, co-culture infection resulted in a significant increase in the apical secretion of IFNγ, IL1-β, IL-6 and IL10 (338±5%, p<0.0001; 161±6%, p<0.05; 593±7% p<0.0001 and 136±3%, p<0.05, respectively) and in the basolateral secretion of IFNγ, IL-1β, IL-4, IL-6 IL10 and TNFα (261±5% p<0.001; 572±4% p<0.0001; 227±5% p<0.0001; 704±18% p<0.0001; 156±8%, p<0.001 and 167±3%, p<0.0001, respectively) (**figure 1b**). The only instance where TNFα was significantly changed in the apical supernatants was in the infected co-cultures when compared to uninfected epithelial cell monocultures with a 329±13%, p<0.01 increase. This data supports a significant augmentation of the inflammatory response to SARS-CoV-2 infection occurs in the presence of pre-existing airway neutrophilia. Importantly, this secretion profile closely reflects the cytokine biomarkers that have been clinically identified in patients hospitalized with severe COVID-19 disease [43–45], highlighting the importance of the co-culture models in recapitulating features associated with more severe responses to SARS-CoV-2 and demonstrating a role for neutrophils in the inflammatory profile observed in patients with severe COVID-19.

### Increased SARS-CoV-2 infection of the airway epithelium is associated with neutrophilia and disruption of epithelial barrier integrity

To determine whether a proinflammatory niche, such as that observed in the presence of pre-existing neutrophilia, impacts epithelial barrier integrity and viral load of the epithelial cells we evaluated barrier resistance and viral content of the airway epithelium. Trans epithelial electrical resistance (TEER) was recorded at 4 and 24 hours after introduction of neutrophils to the airway epithelium. The presence of neutrophils significantly reduced the TEER and, therefore, epithelial barrier integrity, by 23±9%, p<0.05 after 4 hours. This reduction in TEER was sustained through 24 hours (22±4%, p<0.05), all data are compared to epithelial monocultures (**figure 2a**). Evaluation of intracellular viral load by qRT-PCR for SARS-CoV-2 nucleocapsid RNA in the epithelial cells under the same conditions indicated a concurrent and significant increase in infection after the addition of neutrophils by 3.1±1.1-fold (p<0.05) (**figure 2b**). In the absence of infection, no SARS-CoV-2 RNA was detected (data not shown). To determine if the change in epithelial barrier function allowed for increased passage of viral particles from the apical to basolateral surface of the airway epithelium, we also evaluated SARS-CoV-2 spike protein expression in the supernatants **(figure 2c-d).** The presence of neutrophils significantly decreased the apical viral load from 69204±9200.1 fg/ml to 6655.6±475.61 fg/ml (p<0.01) (**figure 2c**) with a concurrent increase in the basolateral viral load from 488.23±129.12 fg/ml to 2307.7±238.94 fg/ml (p<0.01) (**figure 2d**). This data shows that the presence of neutrophils is allows for increased migration of virus from the apical to the basolateral surface. To determine whether the physical presence of neutrophils is essential or whether the pro-inflammatory cytokines released from neutrophils in epithelial co-cultures (**figure 2**) and stimulated by SARS-CoV-2 infection, could induce similar changes in epithelial barrier function, we supplemented the culture media with IFNγ (10 ng/ml), IL-1β (10 ng/ml), IL-6 (10 ng/ml) and TNFα (10 ng/ml) (referred to as cytomix). In the presence of cytomix TEER decreased after 4 hours (18±7%, not significant) with a further and significant decline of 30±5%, p<0.05 after 24 hours (**figure 2e**). This decrease in TEER corresponded to an increase in viral infection of the airway epithelium (2.6±0.5-fold, p<0.05) in the presence of cytomix (**figure 2f**). Reflecting the observations in the presence of neutrophils the apical concentrations of SARS-CoV-2 were decreased from 76703±8708.7 fg/ml to 35261±3598.7 fg/ml (p<0.05) and basolateral concentrations increased from 479.87±129.21 fg/ml to 12344±906.62 fg/ml (p<0.001). This data supports the hypothesis that pro-inflammatory cytokines secreted by neutrophils allow for increased transition of virus from the apical to basolateral surfaces of the airway epithelium.

**Figure 2:**
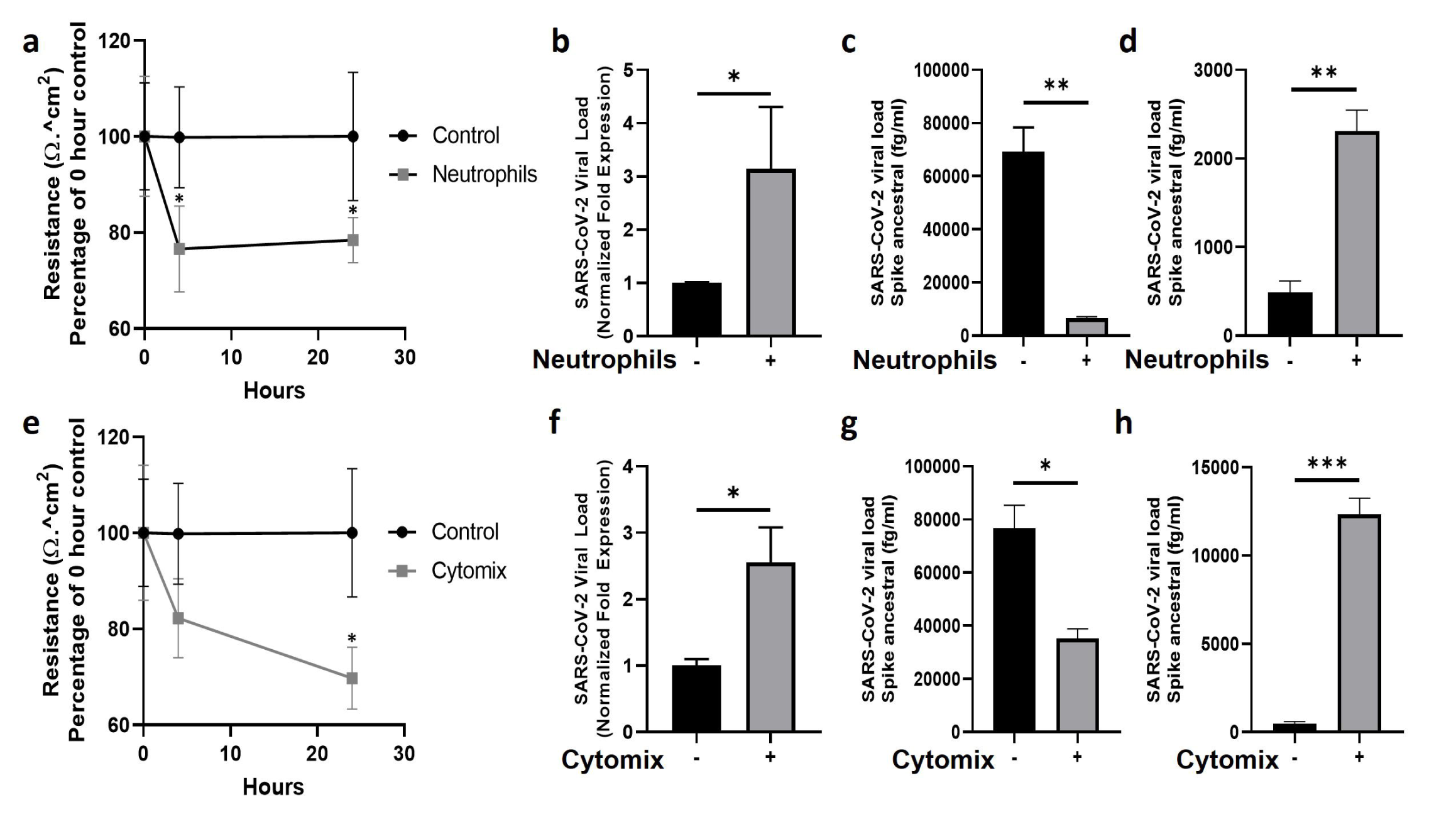
Neutrophils and pro-inflammatory cytokines break down the epithelial barrier and increase viral load in human airway epithelial cells. a) TEER of human airway epithelial cells at the air-liquid interface in the presence, or absence (control), of neutrophils. b) Intracellular viral load of SARS-CoV-2 RNA isolated from infected human airway epithelial cells with neutrophils present. c) Apical supernatant SARS- CoV-2 spike protein concentration 4 hours post infection with neutrophils present. d) Basolateral supernatant SARS-CoV-2 spike protein concentration 4 hours post infection with neutrophils present. e) TEER of human airway epithelial cells cultured with a “cytomix” of TNFα, IL-1β, IL-6 and IFN-γ each at 10ng/ml. f) Intracellular viral load of SARS-CoV-2 in airway epithelial cells cultured with cytomix. g) Apical supernatant SARS-CoV-2 spike protein concentration 4 hours post infection from epithelial cells cultured with cytomix. h) Basolateral supernatant SARS-CoV-2 spike protein concentration 4 hours post infection from epithelial cells cultured with cytomix. Data are expressed as mean±SEM. Statistical significance of TEER data was determined by ANOVA and viral load data was analyzed using an unpaired two-tailed Student’s t-test. *p<0.05. Experiments include n=3 experimental repeats of N=3 independent epithelial donors paired with 3 independent neutrophil donors.

### Neutrophils increase SARS-CoV-2 infection of the epithelium including basal stem cells

To investigate changes in airway pathology associated with SARS-CoV-2 infection we evaluated co- localization of SARS-CoV-2 virus in the presence or absence of neutrophils. Analysis of the airway structure by hematoxylin and eosin (H&E) highlights significant changes in pathology in the presence of neutrophils (**figure 3a-d**). In the absence of neutrophils and infection the airways comprise of a typical airway epithelium with KRT5+ basal cells residing on the basolateral surface and ciliated cells lining the airway lumen (**figure 3a**). Despite the presence of pro-inflammatory cytokines produced by the neutrophils, epithelial cells appear to tolerate the presence of neutrophils, which can be observed in close proximity to the apical ciliated cells in the culture model (**figure 3b)**. In an airway without neutrophils, the epithelial cells are capable of tolerating infection by SARS-CoV-2 after 4 hours of exposure with little evidence of cellular pathology by H&E and only sporadic infection observed in the columnar epithelial cells (**figure 3c and supplemental figure S3)**. Most notably, in the presence of neutrophils, significant cellular pathology is observed by H&E, with evidence for thickening of the basal cell layer, indicative of basal cell proliferation (**figure 3d**). Furthermore, SARS-CoV-2 infection in epithelium is more widespread across the entire epithelial layer with KRT5+ basal cells also being infected (**figure 3d and supplementary figure S3**). In our model system, neutrophils drive significant cellular pathology and increase basal cell proliferation and infection by SARS-CoV-2. Infection of basal cells at such a short timepoint is likely to have significant implications on their function and subsequently airway regeneration.

**Figure 3:**
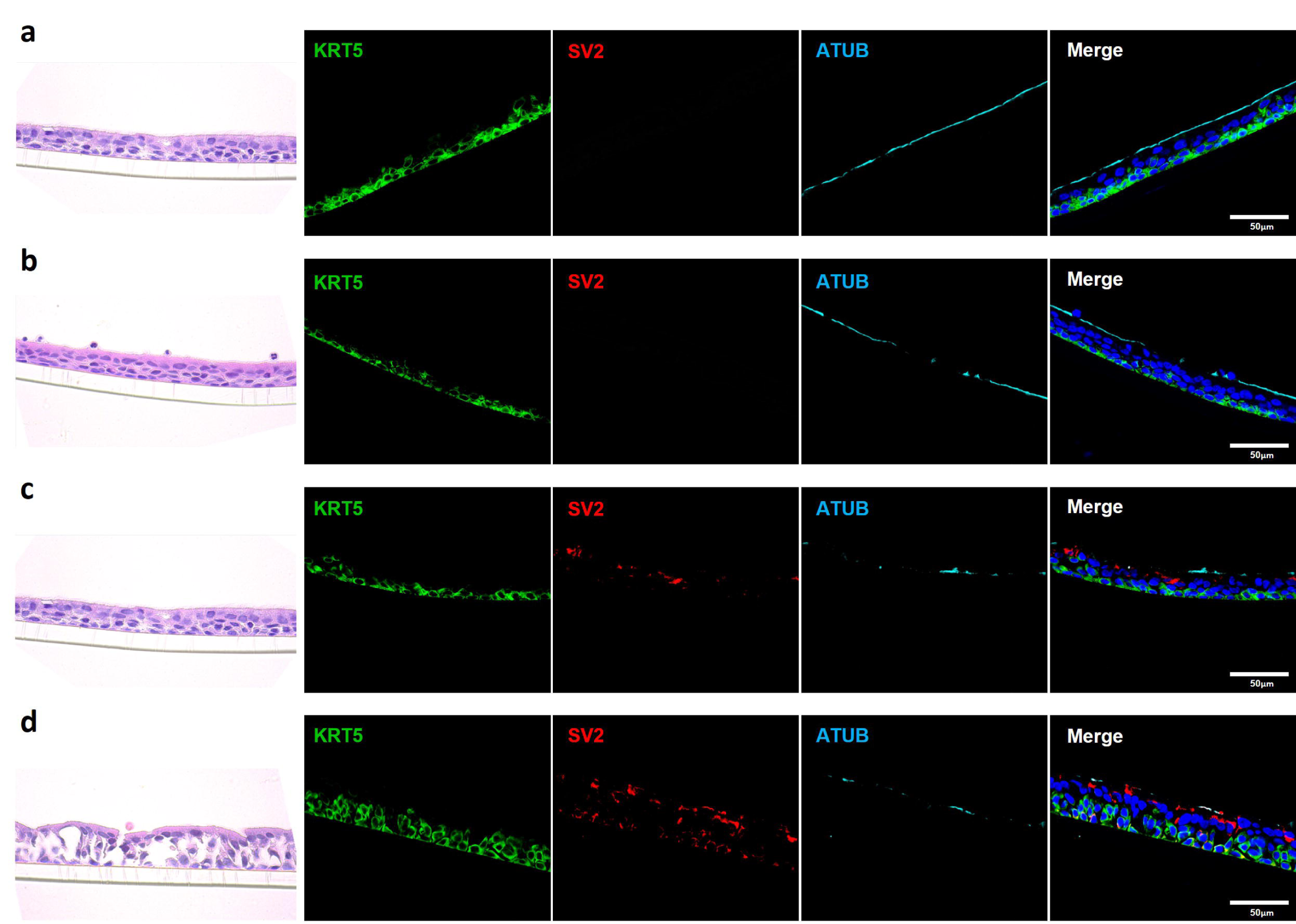
A pre-existing pro-inflammatory environment increases SARS-CoV-2 infection of airway basal stem cells. a-d) representative hematoxylin and eosin (H&E) staining and immunofluorescent images of cross section culture models probed for KRT5 (green) Sars-CoV-2 (red) and alpha-tubulin (cyan). a) uninfected monocultured epithelial cells. b) uninfected epithelial cell – neutrophil co-culture. c) SARS-CoV-2 infected epithelial cell monoculture. d) SARS-CoV-2 infected epithelial cell – neutrophil co-culture. All IF images have nuclei counterstained with DAPI (blue) and scale bars represent 50 μm. All images are representative of 3 independent experimental repeats of 3 neutrophil and 3 epithelial random donor pairings.

### Airway epithelial pathologies are associated with neutrophil activity in severe COVID-19

The data presented from our **in vitro** models suggests that neutrophils play a role in the pathophysiology of early-stage epithelial infection in COVID-19. To further investigate continued neutrophil related pathologies in severe COVID-19 we evaluated epithelial cell related damage and neutrophil activity in post-mortem human tissues from COVID-19 subjects. Formalin-fixed paraffin embedded (FFPE) tissue sections from two post-mortem COVID-19 subjects, kindly provided by the autopsy service at the University of Vermont Medical Center (UVMMC) were assessed for infection-related pathologies through H&E staining. Pathologies were determined by an independent pathologist to be consistent with severe ARDS with mixed inflammatory cell infiltrates, inclusive of neutrophils, and organizing pneumonia (**figure 4a-d**). Tissues from patient Au20-39 (detailed in **supplementary table S1**) contained a mild infiltrate of chronic inflammatory cells surrounding the bronchiole and arterial tissues with involvement in the adjacent surrounding alveolar tissue (**figure 4a** and **supplementary figure S4a**). Scattered giant cells were identified in alveolar spaces and within the interstitium (**figure 4b**, indicated by the red arrows and **supplementary figure S4b**). No well-formed granulomas or definite viral inclusions were evident in this patient. Images from the second patient; Au20-48 **(supplementary table S1)** also show severe organizing diffuse alveolar damage with evidence of barotrauma (**figure 4c** and **supplementary figure S4d**). Alveolar spaces are lined by hyaline membranes or filled with polyps of organizing pneumonia and chronic inflammation (**supplementary figure S4d**). Alveolar walls are expanded with edema and a mixed inflammatory cell infiltrate including neutrophils (**supplementary figure S4c-d**). Bronchioles demonstrate chronic injury with peribronchiolar metaplasia and early squamous metaplasia (**figure 4d** and **supplementary figure S4c**). Organizing pulmonary emboli are present in several arteries (**supplementary figure S4c-d**). There are frequent rounded airspaces lined by inflammatory cells and giant cells, consistent with barotrauma from ventilation injury (**supplementary figure S4d**). There are also scattered giant cells in the interstitium not associated with the barotrauma (**supplementary figure S4c-d**). Given the extensive infiltration of inflammatory cells, inclusive of neutrophils, we further evaluated the neutrophil-related epithelial tissue pathology in both patients. An array of airway tissue pathologies was evident in both tissues including 1) basal cell hyperplasia and small airway occlusion (**figure 4e**), 2) epithelial damage and tissue remodeling of smaller ciliated airways (**figure 4f**), 3) epithelial shedding of large cartilaginous airways (**figure 4g**), 4) neutrophil invasion into the airway lumen (**figure 4h**). and finally, 5) neutrophil invasion in the alveolar space with associated alveolar tissue damage and remodeling (**supplementary figure S4E**). In each of these examples, neutrophils were detected and frequently demonstrated strong neutrophil elastase (NE) activity (**figure 4e-i**), and myeloperoxidase (MPO) expression (a common neutrophil marker) is frequently observed around centers of SARS-CoV-2 infection in postmortem COVID-19 tissues (**figure 4f**). From this data we conclude that neutrophils are a core part of the COVID-19 lung pathophysiology and significantly impact airway infection and injury in response to SARS-CoV-2 infection.

**Figure 4:**
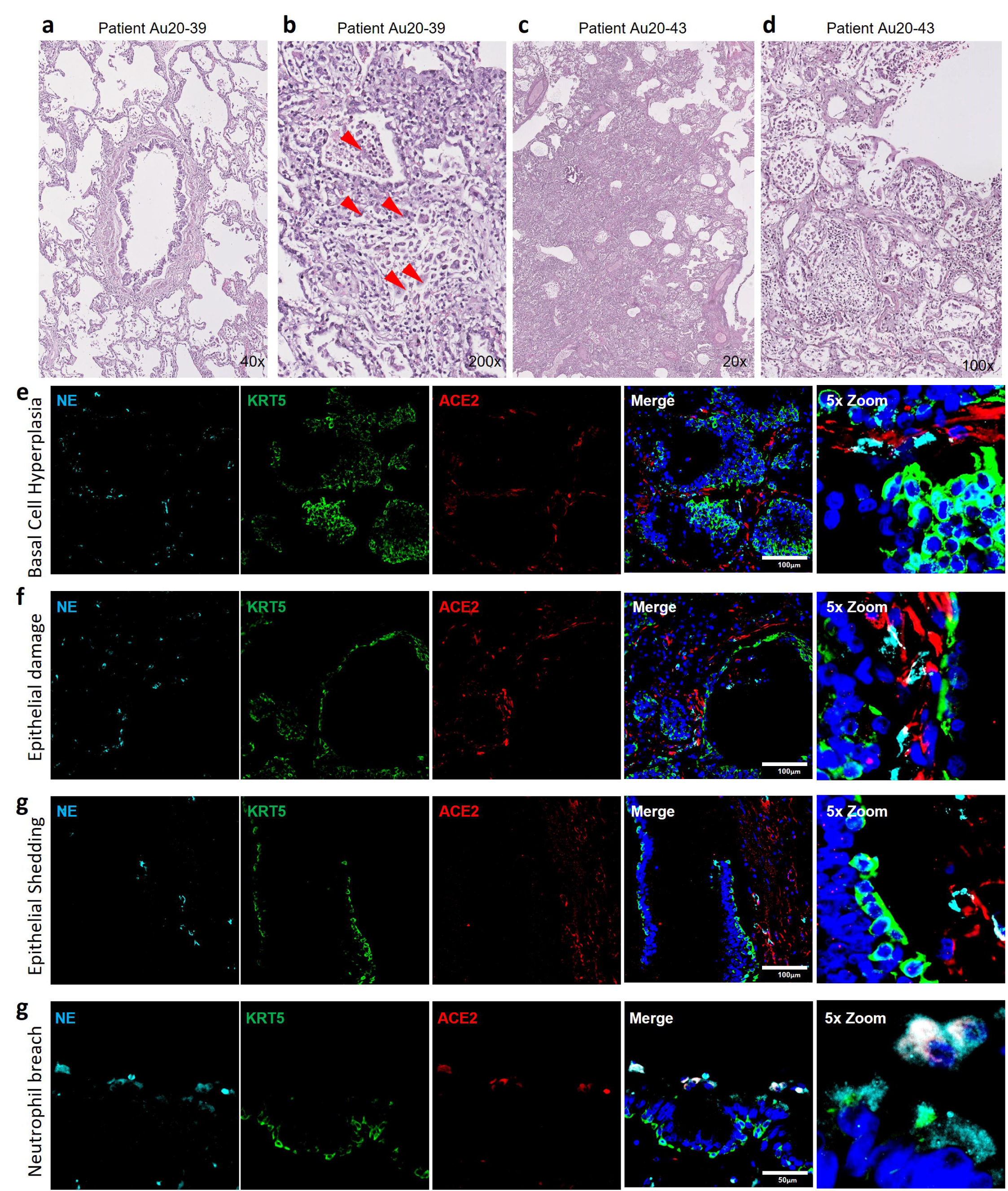
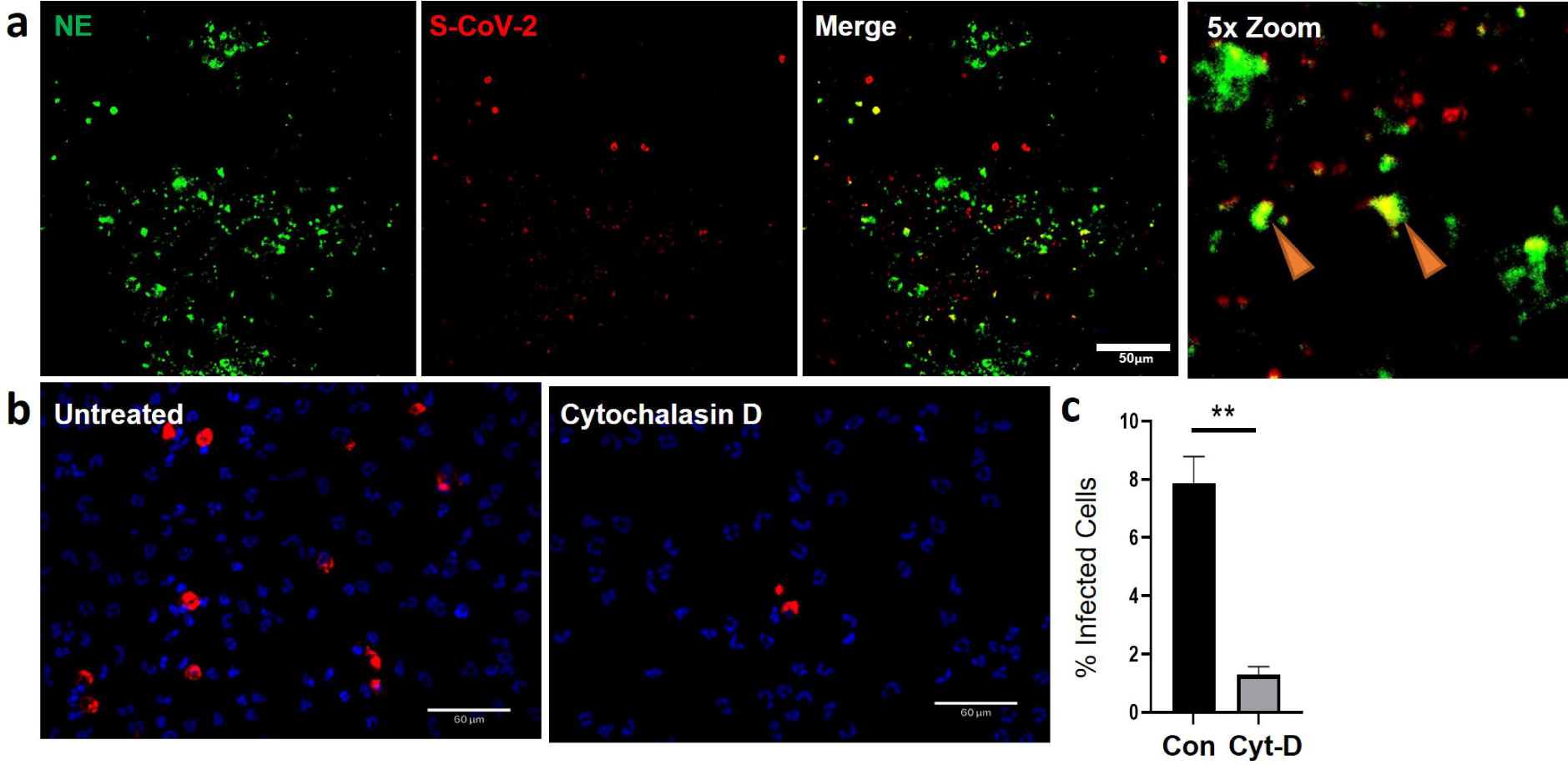
Neutrophil associated tissue pathology in post-mortem COVID19 human lung airways. a-d) Representative images of hematoxylin and eosin (H&E) staining of postmortem COVID-19 patient tissues showing patchy organizing pneumonia centered around a major artery and an airway (a); focally expanded interstitium by a mixed cellular infiltrate including scattered giant cells (red arrowheads) (b); diffuse alveolar damage from intense fibroinflammatory process and barotrauma induced rounded airspaces (c) and organizing diffuse alveolar damage with fibrin disposition replaced by organizing pneumonia, inflammatory cells and oedema (d). e-i) Representative IF images of postmortem COVID-19 tissue probed for NE (cyan), KRT5 (green) and ACE2 (red). Images highlight; small airway occlusion resulting from basal cell hyperplasia with surrounding neutrophils present (e); epithelial damage with breaching neutrophils into the luminal space (f); epithelial shedding, inclusive of basal cell layer with neutrophil inclusion of mucosal surface (g); neutrophil breach into airway luminal space with high neutrophil elastase activity (h) and diffuse neutrophil invasion of alveolar spaces (i). All IF images have nuclei counterstained with DAPI (blue) and scale bars represent 100 μm. All images are representative of 3 independent regions per donor at least 2 independent donors.

### Phagocytosis of SARS-CoV-2 is the predominant mechanism of viral internalization in neutrophils

As previously mentioned, airway diseases, such as CF, that are co-morbidities for severe SARS-CoV-2 infection and progression to severe COVID-19, are also associated with significant infiltration of the airways with neutrophils (**supplementary figure S5a-b**). Interestingly, the neutrophils also colocalized with strong ACE2 expression (**supplementary figure S5).** Despite having significant ACE2 expression our data suggests that internalization of the virus in neutrophils is likely through phagocytosis. The apical concentration of SARS-CoV-2 in the presence of neutrophils was significantly smaller than the apical concentrations of SARS-CoV- 2 in the presence of cytomix **(figure 4c&g)** at 6655.65±475.61 fg/ml compared to 35260.93±3598.7 fg/ml, p<0.01. This suggests that viral clearance is taking place by the neutrophils in their functional role as professional phagocytes. In our experiments SARS-CoV-2 viral RNA was detected in the co-cultures by RNAscope confirming infection of the airway epithelium (**figure 5a**). Interestingly, NE activity was heavily centered around sites of SARS-CoV-2 infection synonymous to that observed in post-mortem patient tissues (**supplementary figure S4f**), and internalization of SARS-CoV-2 by neutrophils was also confirmed by co- localization of staining for NE and SARS-CoV-2 viral RNA (**figure 5a)** *in vitro,* indicated by the orange arrows.

**Figure 5:**
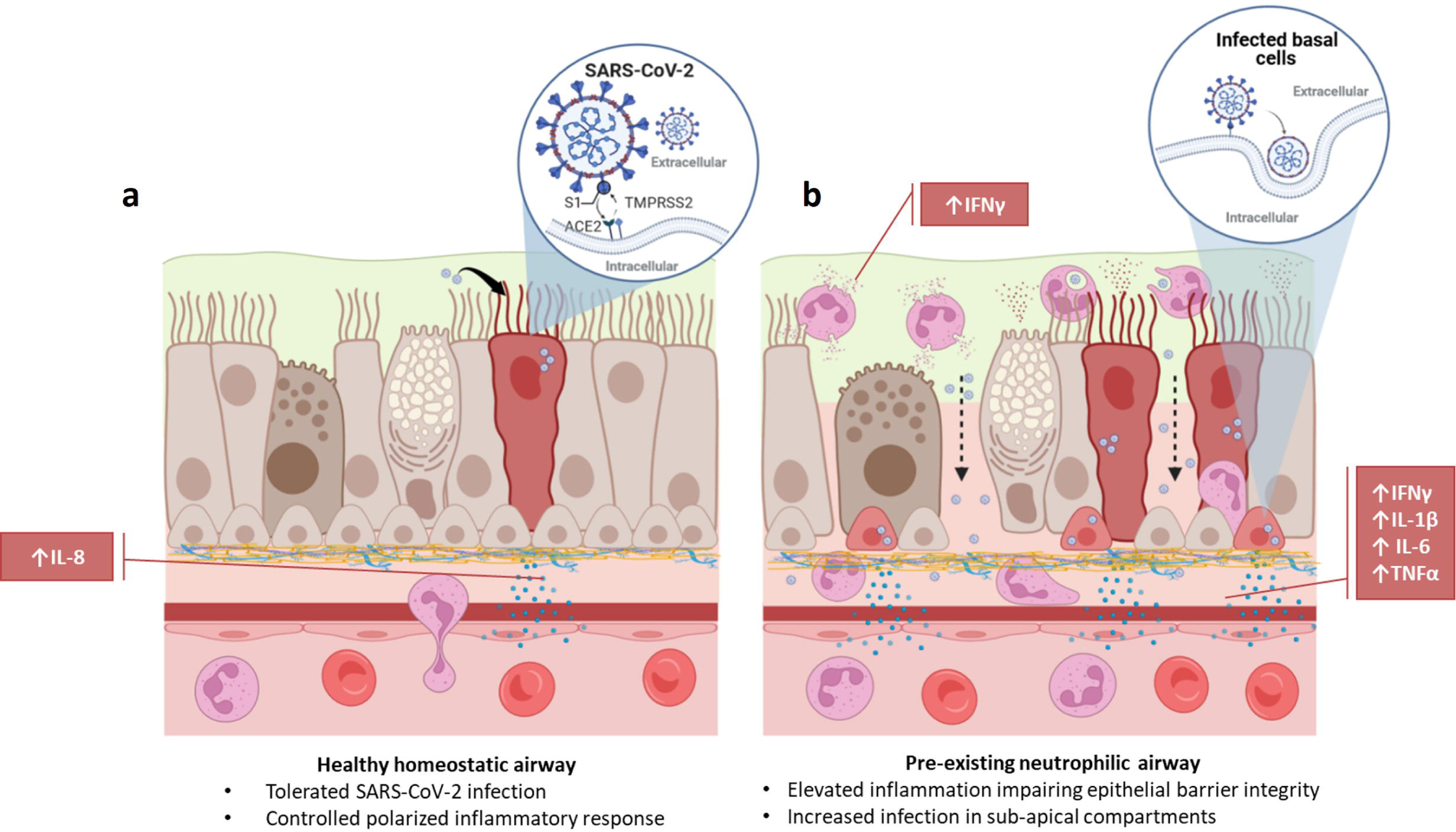
Cytochalasin D inhibits internalization of SARS-CoV-2 in neutrophils. a) Representative IF images of ALI cultures probed for neutrophil elastase (NE) (Green) infected with SARS-CoV-2 (red) detected by RNAScope. b) Quantification or SARS-CoV-2 positive neutrophils relative to total number of neutrophils determined by DAPI (blue). Data expressed as mean±SEM. **p<0.01 unpaired 2-tailed Student’s T-test. N=3 independent neutrophil donors, n=3 experimental replicates.

Finally, to determine whether the expression of ACE2 protein in neutrophils has a significant impact in the response of neutrophils to SARS-CoV-2, we evaluated whether neutrophils were being actively infected via a physical interaction of ACE2 and SARS-CoV-2 or functionally phagocytosing the SARS-CoV-2 virus. The decrease in apical spike protein concentrations when neutrophils are present, compared to epithelial cell monocultures, suggests that the neutrophils are clearing the virus at the apical surface through innate pattern recognition phagocytosis. To better understand this, the frequency of SARS-CoV-2 internalization in monocultures of neutrophils was quantified in the presence or absence of cytochalasin D (15 μM) to inhibit phagocytosis (**figure 5b**). The number of neutrophils positive for SARS-CoV-2 RNA, reflecting viral internalization relative to the total number of neutrophils, was calculated after infection of the cells with SARS-CoV-2 (MOI = 2). Infection, detected by RNA scope, occurred at a rate of 7.9±1% of neutrophils in culture. This signal was significantly reduced by from 7.9±1% to 1.3±0.3% in the presence of cytochalasin D (**Fig. 5C**). Disruption of the actin cytoskeleton, a core component of phagocytosis, therefore, significantly reduced viral uptake in neutrophils. This suggests the primary mechanism for SARS-CoV-2 internalization in neutrophils is phagocytosis.

## Discussion

It is well established that neutrophils are critical in the development of pathological inflammation which can result in both acute and chronic tissue damage. Evaluation of post-mortem COVID-19 tissues indicated significant neutrophil presence and activation in regions of airway epithelial damage and pathology. In addition, we know that many SARS-CoV-2 co-morbidities, including chronic airway disease [30, 46], aging [47–49] and obesity [50–52], are also associated with chronic airway inflammation. In this study we developed a model of pre-existing/chronic airway neutrophilia akin to a model previously developed to investigate other respiratory viruses [27] and applied this to investigate the initial stages of SARS-CoV-2 airway infection. Using this model, we were able to conclude that the pre-existing presence of neutrophils in airway epithelium generates a pro-inflammatory niche, significantly augments initial proinflammatory responses to SARS-CoV- 2 infection, increases viral load in basal stem cells and decreases airway epithelial barrier integrity. Our data, therefore, supports a key role for neutrophilic airway inflammation in determining the infectivity and outcome measures of COVID-19.

Establishing a primary cell co-culture model of an inflammatory airway overcomes some of the limitations of using immortalized cell lines and more complex *in vivo* models. While *in vivo* models are perhaps considered gold standard in infection models, they have not been observed to closely mimic human lung pathophysiology, particularly with respect to SARS-COV-2. While infection can be detected, no animal model had closely reflected COVID-19 pathogenesis that leads to severe symptoms and fatal lung disease [5, 53]. Furthermore, studying neutrophilia in animal models is challenging, several depleted or knockout models exist [54], however evaluation of elevated lung neutrophilia typically requires pro-inflammatory stimulation with lipopolysaccharide (LPS) [55], this could complicate interpretation of findings in relation to viral infection. Our models use primary HBECs, some of the first cells exposed to the virus that express endogenous levels of ACE2 and TMPRSS2. This allowed for investigation of the initial stages of SARS-CoV-2 infection and characterization of acute phase inflammatory responses.

Neutrophil phenotype and function, including those involved in resolving viral infections, is strongly regulated by signals received from their tissue micro-environment [56], in our study we considered neutrophil responses in the presence of an epithelial micro-environment. Our model mimics components of neutrophilic airway inflammation associated with other chronic lung diseases that have been linked with a predisposition to developing more severe COVID-19 disease. Perhaps our most striking finding is the presence of a differential polarized inflammatory response in response to neutrophils and/or SARS-CoV-2. IL-8, the core chemoattractant for neutrophils [38–40, 57], is secreted only on the basolateral surface of the epithelial monocultures, demonstrates that epithelial cells are capable of recognizing neutrophils within their niche and downregulate this chemokine secretion as a result and that the model recapitulates the directionality required to recruit circulating neutrophils into an infected epithelial environment. Furthermore, despite seeding neutrophils on the apical surface of our model, we observed a predominant pro-inflammatory niche basolaterally, with increases in IL-1β, IL-4, IL-6 and TNFα. Through paired comparisons to primary airway epithelial cells in monoculture, we were able to demonstrate key differences in the secretion of pro- (IFNγ, IL1β, IL-6, IL-8 and TNFα) and anti-inflammatory (IL-4 and IL10) mediators, epithelial barrier integrity and infectivity of epithelial cells (**figures 1-2**), which would have been over-looked in monoculture experiments involving airway infection only. Importantly, the secretion of pro-inflammatory cytokines in our model is consistent with clinical studies that have reported an elevated inflammatory profile associated with severe COVID-19 disease. In patient peripheral blood samples, IL-6 [58–61] IL-10 [59, 60] are consistently higher in COVID-19 patients and correlate with disease severity. Additionally, IL-6 and IL-8 are even higher in ICU than the IMU [62]. Our data also closely mimics responses observed in primate models of the disease [63]. The lack of robust inflammatory response of the epithelium alone may also provide rational for why some people are predisposed to more severe responses than others. In fact, our data evaluating the response of the more proximal, cartilaginous airways may highlight the importance of a robust proximal airway defense mechanism that controls the progression to severe COVID-19 associated with ARDS and distal airway dysfunction.

Pro-inflammatory cytokines, including IFNγ, IL1β, IL-6 and TNFα, have extensively been shown to disrupt barrier integrity and permeability of the epithelium [64, 65]. This breakdown in barrier integrity exists to allow for leukocyte migration to sites of stress and infection. Theoretically, any tight-junction breakdown that allows for more leukocyte migration, would also allow for increased permeability for viral particles to sub-apical and sub-epithelial structures, thus increasing infectivity and cellular viral loads. Our data supports this phenomenon with both neutrophils and cytomix synonymously decreasing barrier integrity **(figure 2)** whilst increasing intracellular viral loads and virus concentrations in sub-apical compartments. This association of epithelial barrier integrity with an increase in intracellular epithelial viral loads, especially in the basal stem cells, suggests that epithelial barrier integrity plays an important functional role in SARS-CoV-2 infection.

Finally, we addressed the key question of whether neutrophils, as professional phagocytes [66, 67], are capable of innate recognition of SARS-CoV-2 as an invading pathogen through innate recognition pathways, and/or are capable of infection by SARS-CoV-2 inherently via ACE2 expression. Our data supports a high level of expression of ACE2 at the protein level, but not the RNA level in neutrophils; an observation recently reported by Veras and colleagues [26]. Furthermore, infection is facilitated by TMPRSS2 and we did not see any evidence for expression on neutrophils. By using cytochalasin D to breakdown actin filament organization we significantly reduced internalization, supporting a predominant role for phagocytosis in the internalization of SARS-CoV-2 in neutrophils. Reports are, however, emerging that suggest a significant role for cytoskeletal rearrangement in SARS-CoV-2 entry and, therefore, we cannot entirely rule out infection [68]. The use of blocking antibodies has potential to elucidate the mechanisms of internalization, however, neutrophils express copious amounts of Fc receptors [69] and likely to recognize antigens and opsonize through phagocytosis. Our assay attempted to investigate an innate recognition, i.e. a non-humoral opsonization of the SARS-CoV- 2 virus. To determine whether the expression of ACE2 on neutrophils is functionally relevant in SARS-CoZV- 2 infection further investigation will be essential.

In conclusion, we have developed a model to study neutrophil-epithelial interactions which more closely reflects an *in vivo* and more clinically relevant infection of airways than monocultures. Our findings demonstrate that the co-presence of neutrophils generates a polarized pro-inflammatory niche with the conducting airway epithelium that is significantly augmented with SARS-CoV-2 infection. This pro- inflammatory niche breaks down the epithelial barrier integrity allowing for increased epithelial infection including basal stem cells. Overall, this study reveals a key role for pre-existing chronic airway neutrophilia in determining infectivity and outcomes in response to SARS-CoV-2 infection that highlight neutrophilia as a potential target for prevention of severe COVID-19 disease.

## Acknowledgements

The authors acknowledge the Center for Advanced Research Computing (CARC) at the University of Southern California for providing computing resources that have contributed to the research results reported within this publication. URL: https://carc.usc.edu. All Biosafety Level 3 work was performed within The Hastings Foundation and The Wright Foundation Laboratories at USC. SARS-V2 BSL3 resources supported by a grant from the COVID-19 Keck Research Fund. Flow cytometry was performed in the USC Flow and Dr. Scott Randell at the University of North Carolina Marsico Lung Institute Tissue Procurement and Cell Culture Core supported in part by NIH DK065988 and grant BOUCHR19R0 from the Cystic Fibrosis Foundation, for their partnership in providing lung tissues for research. Finally, the authors wish to thank the USC COVID-19 Biospecimen Repository for assistance with pathologic analysis

## Author Contributions

Conceptualization, B.A.C and A.L.R.; Methodology; B.A.C and A.L.R.; Formal Analysis, M.P.S.; Investigation, B.A.C, E.J.Q, Z.L., N.D., C.N.S., S.K., W.D.W., J.H., and A.L.R; Writing - Original Draft, B.A.C and A.L.R., Writing – Review and editing, B.A.C and A.L.R., Funding Acquisition, A.L.R.

