## Supplemental Information for "Neutrophilic inflammation promotes SARS-CoV-2 infectivity and augments the inflammatory responses in airway epithelial cells"

**Supplemental Methods**

***In vitro* airway differentiation: Spheroids**

For differentiated spheroids, transwells were pre-treated with 25% (v/v) of Matrigel in Pneumacult Ex+ (Stem Cell Technologies) for at least 1 hour at 37°C. For spheroid formation, a single cell suspension of HBECs was generated using with Accutase. Single cells were resuspended in Matrigel™ 5% in Pneumacult Ex+. The cell suspension was loaded onto the pre-established gel-bed and Pneumacult Ex+ media added to the lower chamber. After 3 days the media was exchanged for Pneumacult ALI maintenance media (Stem Cell Technologies) and changed every 2~3 days.  Spheroids were fixed with 4% PFA in PBS at 5, 10, 14 and 20 days after the culture. For immunostaining, spheroids, attached at the bottom of the transwells, were pre-treated with 0.1% of Triton X-100 in PBS (hereinafter 0.1% PBT) for 10 min for permeabilization. The cells were incubated with primary antibody for 1hr 30min at room temperature and washed with 0.1% PBT. Secondary antibodies were incubated for 45 min at room temperature and washed with 0.1% PBT. Nuclei were stained with DAPI. The spheroids were mounted with Fluoromount G (#0100-01; SouthernBiotech). The spheroid images were taken with confocal microscope (Zeiss, LSM710) using a Plan Apochromat 63x/1.4 NA oil-immersion objective, ~10 μm of thickness with 0.4μm of z-stack interval and processed with ZEN software (Zeiss) and ImageJ software.

**Single Cell RNA sequencing and Bioinformatic Analysis**

Gene expression counts (GEO accession number: GSE150674, [1]) from both the fresh isolates and ALI cultures were processed using functions in the R package Seurat (version 4.0.2) [2]. For each sample, low quality cells and potential cell doublets were removed by filtering cells based on the number of genes detected and the fraction of reads mapping to the mitochondrial genome. Cells with less than 200 genes detected or greater than 4,900 – 7,000 genes detected (depending on the sample) were removed as well as cells with greater than 10% of mitochondrial reads. The filtered gene counts were then integrated into a unified data set using the

SCTransform [3] based integration workflow implemented in Seurat. The first 30 PCs were used as the number of dimensions for the FindNeighbors and FindClusters functions. To find the optimal clustering resolution value we explored a range of resolution values from 0 to 2 by increments of 0.2. We then visualized the range of cluster resolutions using the R package clustree (version 0.4.3) [4]. A final resolution of 0.4 was selected and used for downstream analyses. Clusters were visualized in two-dimensional space using the RunUMAP function in Seurat. Cluster specific cell types were annotated manually by examining the expression of canonical marker genes within a given cluster and clusters with shared marker gene expression patterns were collapsed. The fresh isolates and ALI couture data sets were analyzed separately.

**Supplemental Results**

Additions to the main text references to the supplemental data

**Expression of ACE2 and TMPRSS2 in human lung and *in vitro* human lung models.** To validate the expression patterns, we re-analyzed published single cell sequencing datasets (GEO accession number: GSE150674) on proximal human lung tissues and found that ACE transcripts were low in abundance and detectible levels of both ACE2 and TMPRSS2 were found in ciliated cells, SMG cells, basal cells, and secretory cells (**supplementary figure S2a-b**). At the protein level expression of ACE2 was observed most notably in ciliated columnar epithelial cells (**supplementary figure S2c**) and in submucosal glands (**supplementary figure S2e**). TMPRSS2 was found to be more ubiquitous in its expression and found throughout the tissue (**supplementary figure S2d&f**) inclusive of strong expression in the ciliated epithelium (**supplementary figure S2d**) and in submucosal glands (**supplementary figure S2f**).

**Expression of ACE2 in infiltrating neutrophils in human lung and *in vitro* human lung models.** The presence of a very high ACE2 expressing cell (**supplementary figure S5a)** with a polynuclear migratory phenotype was noted to be located in blood vessels, tissue parenchyma and infiltrating the epithelium. This cell type was substantially more prevalent in CF tissues (**supplementary figure S5b)** compared to patients with no underlying chronic respiratory disease (**supplementary figure S5a)** and co-localized with Neutrophil Elastase (NE) **(supplementary figure S5c)** and CD15 (**supplementary figure S5d**), confirming the cells as neutrophils.

To confirm neutrophil expression of ACE2 CD15+ peripheral blood polynucleocytes were isolated and co-stained with NE and ACE2 (**supplementary figure S5e**).

### Supplementary Tables

**Supplementary table S1**: Demographic information for lung tissue donors used in this project.

| Identifier | Disease | Age | Gender | BMI | Co-morbidities | Source |
|---|---|---|---|---|---|---|
| Au20-39 | COVID-19 | 39 | Male | 53.2 $Kg/m^2$ | None reported | UVM |
| Au20-48 | COVID-19 | 72 | Male | 29.1 $Kg/m^2$ | GI Reflux, Prostate CA s/p radiation, hyperlipidemia, former smoker, hypertension | UVM |
| 004-H | No-underlying disease | UK | UK | UK | UK | UI |
| 016-CF | Cystic Fibrosis (ΔF508/I633K) | 54 | Female | N/A | Bronchiectasis | UI |
| 018-H | No-underlying disease | UK | UK | UK | UK | UI |
| 019-CF | Cystic Fibrosis (ΔF508/G551D) | 24 | Female | N/A | None reported | UI |

UK, unknown, UI, University of Iowa, UVM, University of Vermont

**Supplementary table S2**: Demographic information for primary human bronchial epithelial cell donors and pairings with blood donors for ALI co-cultures.

| Identifier | Disease | Age | Gender | BMI | Co-morbidities | Source | Blood Donor Pairing |
|---|---|---|---|---|---|---|---|
| 044-H | None known | 73 | Female | 24.8 $Kg/m^2$ | Seizures, Epilepsy, early signs of dementia | IIAM | BD001 |
| 033-H | None known | UK | UK | UK | UK | UI | BD002 |
| 014-H | None known | 24 | Male | UK | UK | UNC-Chapel Hill | BD002 |
| 032-H | None known | UK | UK | UK | UK | UI | BD003 |
| 015-H | None known | UK | UK | UK | UK | UI | BD003 |
| 017-H | None known | 35 | Female | UK | UK | UNC-Chapel Hill | BD004 |

UK, unknown, UI, University of Iowa, UNC-Chapel Hill, University of North Carolina-Chapel Hill

**Supplementary table S3**: Demographic information for primary human blood donors

| Identifier | Disease Sate | Age | Gender |
|---|---|---|---|
| BD001 | No underlying disease | 40 | Female |
| BD002 | No underlying disease | 33 | Male |
| BD003 | No underlying disease | 23 | Female |
| BD004 | No underlying disease | 28 | Female |

**Supplementary table S4**: Key resources table.

| REAGENT or RESOURCE | SOURCE | IDENTIFIER |
|---|---|---|
| ***Antibodies*** | | |
| Chicken Polyclonal anti Cytokeratin 5 (KRT5) | Biolegend | Cat# 905901; RRID:AB_2565054 |
| Rat monoclonal anti Angiotensin Converting Enzyme – 2 (ACE2) | Biolegend | Cat# A20069I; RRID:AB_2860959 |
| Rabbit polyclonal anti Angiotensin Converting Enzyme – 2 (ACE2) | Abcam | Cat# ab15348; RRID:AB_301861 |
| Goat polyclonal anti Epithelial Cell Adhesion Molecule (EpCAM) | R&D Systems | Cat# AF960; AB_355745 |
| Mouse monoclonal anti α-Tubulin (ATUB) | Abcam | Cat# ab24610; RRID:AB_448182 |
| Rabbit monoclonal anti Transmembrane Protease Serine 2 (TMPRSS2) | Abcam | Cat# ab109131 RRID:AB_10863728 |
| Rabbit monoclonal anti Neutrophil Elastase (NE) | Thermo Fisher Scientific | Cat# MA5-32548; RRID:AB_2809825 |
| Mouse monoclonal anti Cluster of Differentiation 15 (CD15) | Thermo Fisher Scientific | Cat# MA5-17042; RRID:AB_2538514 |
| Mouse monoclonal anti Cluster of Differentiation 15 (CD15) – PE Conjugated | Biolegend | Cat# 323005 RRID:AB_756011 |
| Rabbit polyclonal anti SARS-CoV-2 spike glycoprotein | Abcam | CAT# ab272504 RRID: AB_2847845 |
| RNAscope Probe - V-nCoV2019-S | ACDBio | Cat# 848561 |
| ***Bacterial and Virus Strains*** | | |
| SARS-CoV-2 Isolate USA-WA1/2020 | BEI Resources | Cat# NR-52281 |
| ***Biological Samples*** | | |
| AU20-039 Post-mortem COVID human lung tissue | University of Vermont | N/A |
| AU20-043 Post-mortem COVID human lung tissue | University of Vermont | N/A |
| 004-H Post-mortem no underlying respiratory disease lung tissue | Laboratory of Amy Ryan | N/A |
| 016-CF Explanted transplant Cystic Fibrosis Human lung tissue | Laboratory of Amy Ryan | N/A |
| 018-H Post-mortem no underlying respiratory disease lung tissue | Laboratory of Amy Ryan | N/A |
| 019-CF Explanted transplant Cystic Fibrosis Human lung tissue | Laboratory of Amy Ryan | N/A |
| ***Chemicals, Peptides, and Recombinant Proteins*** | | |
| Triton-X | Sigma Aldrich | Cat# 9002-93-1 |
| Paraformaldehyde | Thermo Fisher Scientific | Cat# 28908 |
| Human recombinant Tumor Necrosis Factor-Alpha Research Grade (TNFα) | Miltenyi Biotec | Cat# 130-094-015 |

| | | |
|---|---|---|
| Human Recombinant Interleukin 6 (IL6) | Miltenyi Biotec | Cat# 130-095-365 |
| Human Recombinant Interleukin 1-beta (IL1β) | Miltenyi Biotec | Cat# 130-093-895 |
| Human Recombinant Interferon-gamma (IFNγ) | PeproTech | Cat# 300-02-100ug |
| Flash Phalloidin Red 594 | Biolegend | Cat# 424203 |
| Cytochalasin D | VWR | Cat# C8273-1MG |
| Trizol™ | Thermo Fisher Scientific | Cat# 15596026 |
| Pneumacult-EX Plus Medium | Stem Cell Technologies | Cat# 05040 |
| Pneumacult-ALI Medium | Stem Cell Technologies | Cat# 05001 |
| Hanks balanced Salt Solution | Invitrogen | Cat# 14025076 |
| B-ALI Growth Medium | Lonza | Cat# 00193516 |
| Phosphate Buffered Saline | Invitrogen | Cat# 14190-250 |
| Foetal Bovine Serum | Fisher Scientific | Cat# 16-141-061 |
| Ethylenediaminetetraacetic acid | Thermo Fisher Scientific | Cat# AM9261 |
| TruStain Fc receptor blocker | Biolegend | Cat# 422302 |
| Airway Epithelial Growth Medium | Promocell | Cat# C-21060 |
| Accutase | Stem Cell Technologies | Cat# 07920 |
| Dulbecco's Modified Eagles Medium | Thermo Fisher Scientific | Cat# 11995073 |
| Puromycin | Thermo Fisher Scientific | Cat# A1113802 |
| OptiMEM | Thermo Fisher Scientific | Cat# 31985062 |
| **Critical Commercial Assays** | | |
| MSD Cytokine Assay V-plex Viral Panel 1 Human Kit | MSD | Cat# K15345D-1 |
| MSD SARS-CoV-2 Spike S-PLEX | MSD | Cat#K150ADJS |
| QuantiTect Virus Kit | Qiagen | Cat# 211011 |
| Direct-zol RNA Microprep kit | Zymo | Cat# R2061 |
| SARS-CoV-CDC RUO primers and probes | Integrated DNA Technologies | Cat# 1000673 |
| EasySep direct neutrophil isolation kit | Stem Cell Technologies | Cat# 19666 |
| Hematoxylin | VWR | Cat# 10143-612 |
| Eosin | VWR | Cat# 10143-132 |
| Bovine Serum Albumin | RPI | Cat# A30075-100.0X |
| Normal Donkey Serum | Jackson Immuno Research | Cat# 017-000-121 |
| Paraformaldehyde | Thermo Fisher Scientific | Cat# 28908 |
| Fluoromount-G | SouthernBiotech | Cat# 0100-01 |
| 4′,6-diamidino-2-phenylindole | Thermo Fisher Scientific | Cat# D1306 |
| ***Experimental Models: Cell Lines*** | | |
| Human Basal Epithelial Cells (HBECs) 032H | Laboratory of Amy Ryan | N/A |
| Human Basal Epithelial Cells (HBECs) 015H | Laboratory of Amy Ryan | N/A |
| Human Basal Epithelial Cells (HBECs) 017H | Laboratory of Amy Ryan | N/A |
| Human Basal Epithelial Cells (HBECs) 044H | Laboratory of Amy Ryan | N/A |

| | | |
|---|---|---|
| Human Basal Epithelial Cells (HBECs) 033H | Laboratory of Amy Ryan | N/A |
| Human Basal Epithelial Cells (HBECs) 014H | Laboratory of Amy Ryan | N/A |
| Human CD15+ Neutrophils BD001 | Laboratory of Amy Ryan | N/A |
| Human CD15+ Neutrophils BD002 | Laboratory of Amy Ryan | N/A |
| Human CD15+ Neutrophils BD003 | Laboratory of Amy Ryan | N/A |
| Human CD15+ Neutrophils BD004 | Laboratory of Amy Ryan | N/A |
| VeroE6-hACE2 | Laboratory of Dr. Jae Jung, USC | |
| ***Oligonucleotides*** | | |
| 2019-nCoV_N1-F 2019-nCoV_N1 Forward Primer GAC CCC AAA ATC AGC GAA AT | Integrated DNA Technologies | Cat# 1000673 |
| 2019-nCoV_N1-R 2019-nCoV_N1 Reverse Primer TCT GGT TAC TGC CAG TTG AAT CTG | Integrated DNA Technologies | Cat# 1000673 |
| 2019-nCoV_N1-P 2019-nCoV_N1 Probe FAM-ACC CCG CAT TAC GTT TGG TGG ACC-BHQ1 FAM, BHQ-1 125nM | Integrated DNA Technologies | Cat# 1000673 |
| RP-F RNAse P Forward Primer AGA TTT GGA CCT GCG AGC G | Integrated DNA Technologies | Cat# 1000673 |
| RP-R RNAse P Reverse Primer GAG CGG CTG TCT CCA CAA GT | Integrated DNA Technologies | Cat# 1000673 |
| RP-P RNAse P Probe: FAM – TTC TGA CCT GAA GGC TCT GCG CG – BHQ-1 | Integrated DNA Technologies | Cat# 1000673 |
| ***Software and Algorithms*** | | |
| Image J | Fiji – NIH | |
| Graph Pad Prism | GraphPad | |
| BD *FACSDiva*™ | BD Biosciences | |
| FlowJo | BD Biosciences | |
| R Package Seurat | R Foundation | |
| R Package Clustree | R Foundation | |
| Leica LAS X Navigator | Leica | |
| Discovery Workbench | MSD | |
| ZEN lite | Zeiss | |

### Supplementary figures and figure legends

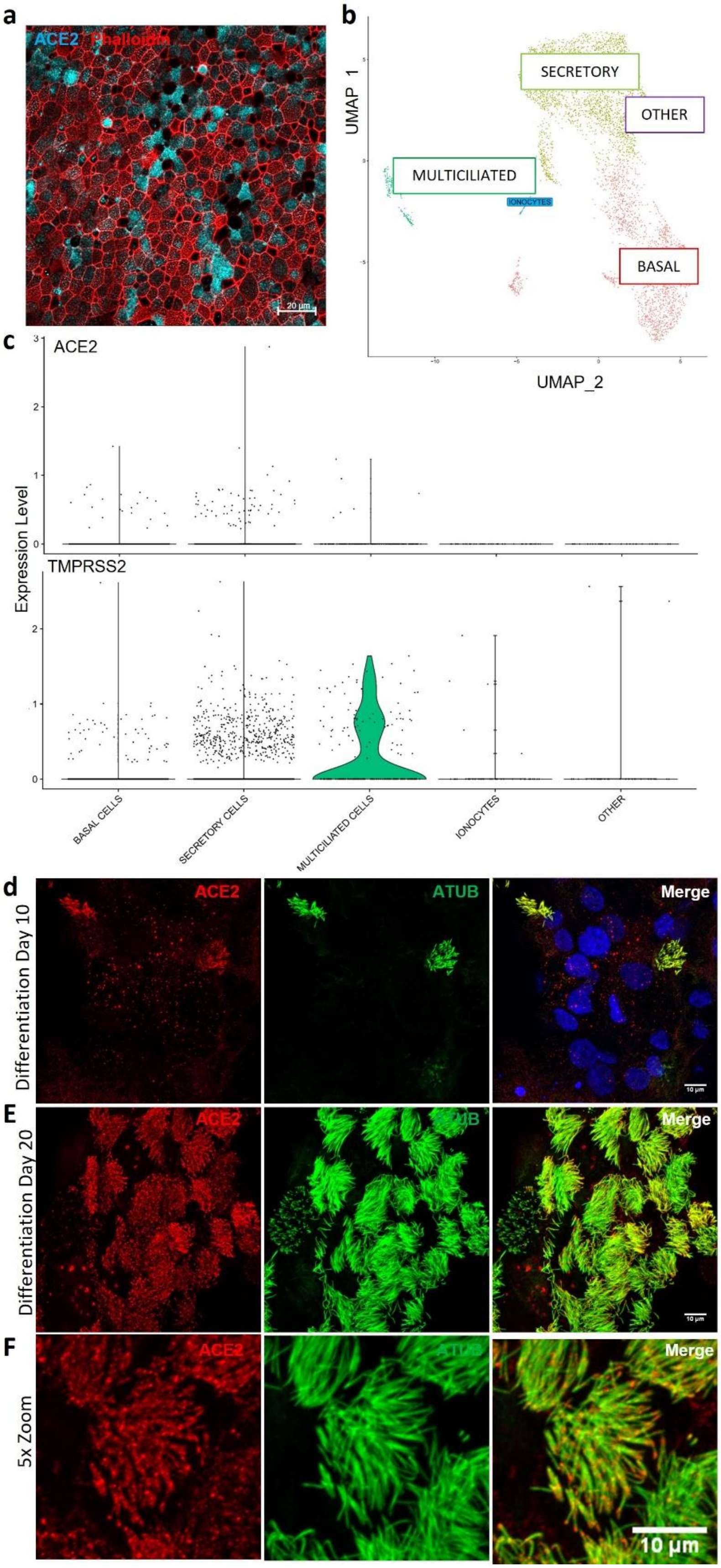

**Supplementary figure S1, related to figure 1: ACE2 expression in *in vitro* ciliated cell basal bodies and axonemes.** a) Representative immunofluorescent image of mature airway epithelium differentiated at the Air-Liquid-Interface probed for ACE2 (Cyan) and co-stained for tight junctions with phalloidin (Red). b) Single cell sequencing cluster analysis visualized by Uniform Manifold Approximation and Projection (UMAP) of cells differentiated at the Air-Liquid-Interface. c) Expression analysis of ACE2 (top panel) and TMPRSS2 (bottom panel) of differential clusters identified UMAP analysis of single cells isolated from differentiated cultures at the Air-Liquid-Interface. D-f) representative immunofluorescent images of tracheospheres probed for ACE2 (red) and acetylated-alpha tubulin (Green) depicting an increase in expression from day 10 differentiation (d) to day 20 differentiation (e) with co-localization of ACE2 at the basal bodies of the cilia (f). Images have nuclei counterstained with DAPI and scale bars are 20 μm in A or 10 μm in d-f.

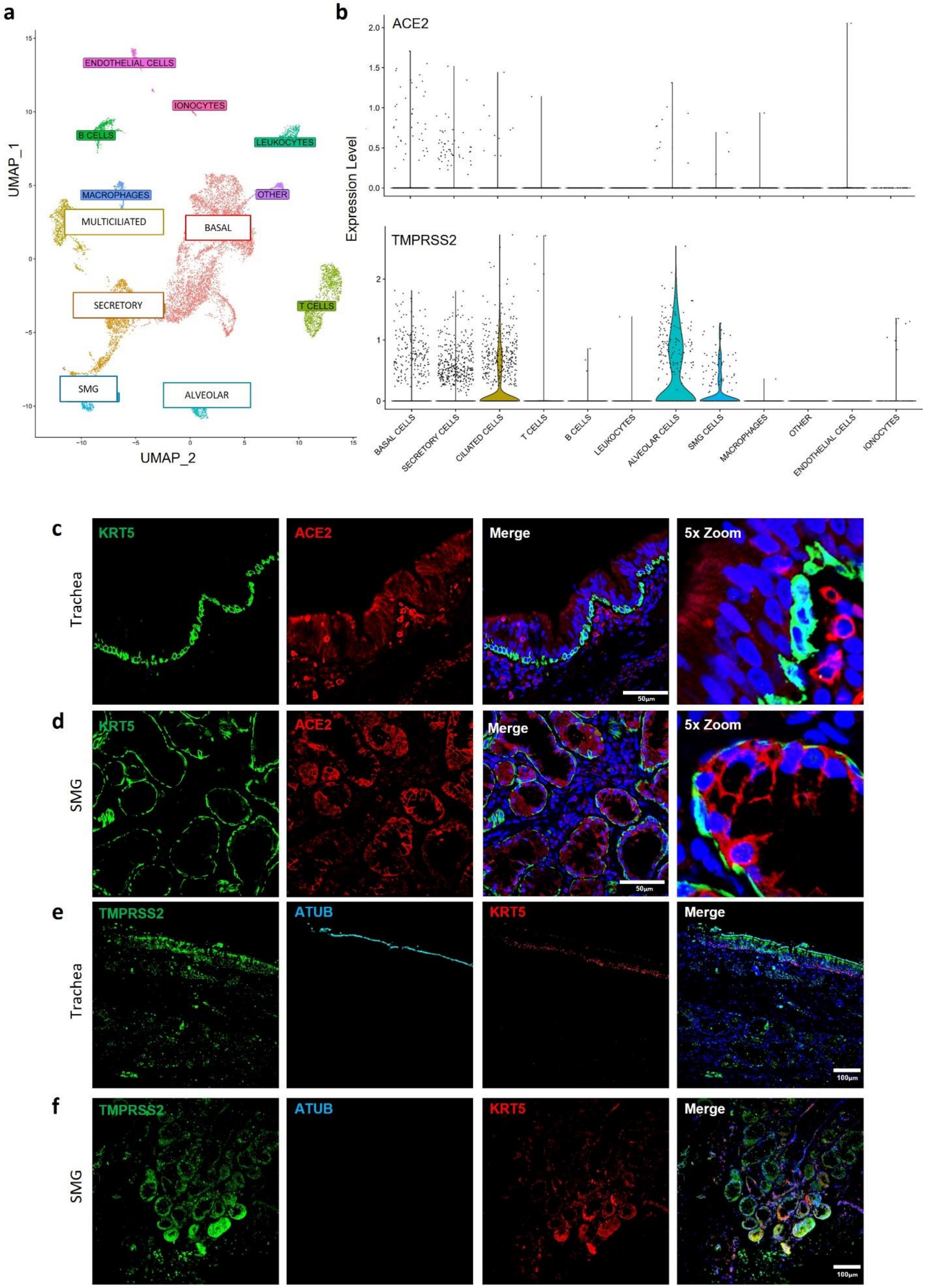

**Supplementary figure S2, related to figure 1: ACE2 and TMPRSS2 expression profiles across human lung upper airways**: a) Single cell sequencing cluster analysis visualized by Uniform Manifold Approximation and Projection (UMAP) of cells isolated from primary human lung tissue. b) Expression analysis of ACE2 (top panel) and TMPRSS2 (bottom panel) of differential clusters identified UMAP analysis of single cells isolated from human lung tissue. Representative immunofluorescent (IF) images of: c) primary human lung trachea surface airway epithelium and d) submucosal glands (SMG) stained for cytokeratin 5 (KRT5, green) and ACE2 (red); e) primary human lung trachea surface airway epithelium and f) submucosal glands (SMG) stained for TMPRSS2 (green), acetylated α tubulin (ATUB, cyan) and KRT5 (red). All images have nuclei counterstained with DAPI (blue) and scale bars represent 50 μm in C and D and 100 μm in e and f. All images are representative of 3 of 3 independent regions per donor at least 2 independent donors

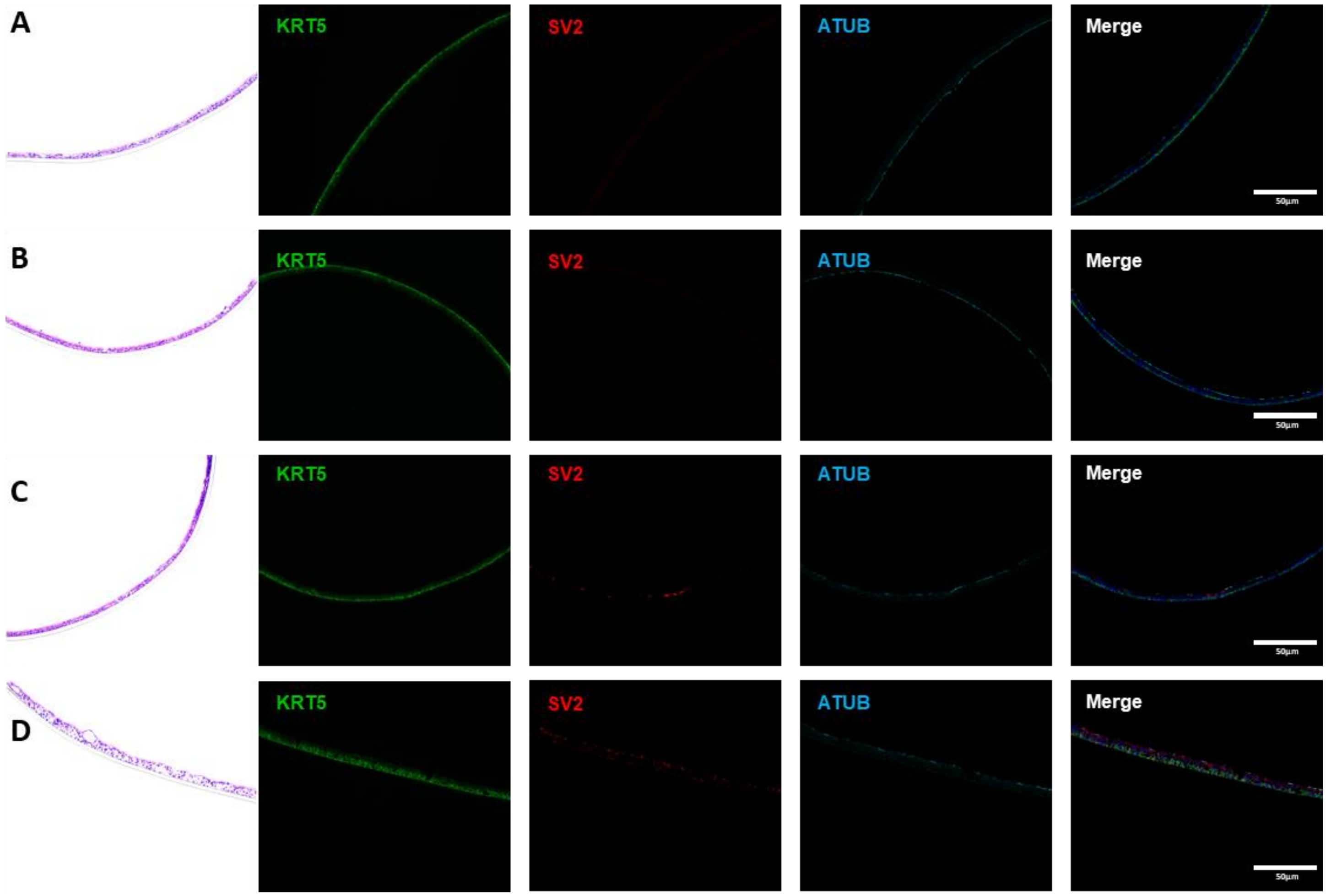

**Supplementary figure S3, related to figure 3**: **A pre-existing pro-inflammatory environment increases SARS-CoV-2 infection of airway basal stem cells**. a-d) representative hematoxylin and eosin (H&E) staining and immunofluorescent images of cross section culture models probed for KRT5 (green) SARS-CoV-2 (red) and alpha-tubulin (cyan) at 10x zoom. a) uninfected monocultured epithelial cells. b) uninfected epithelial cell – neutrophil co-culture. c) SARS-CoV-2 infected epithelial cell monoculture. d) SARS-CoV-2 infected epithelial cell – neutrophil co-culture. All IF images have nuclei counterstained with DAPI (blue) and scale bars represent 50µm. All images are representative of 3 independent experimental repeats of 3 neutrophil and 3 epithelial random donor pairings.

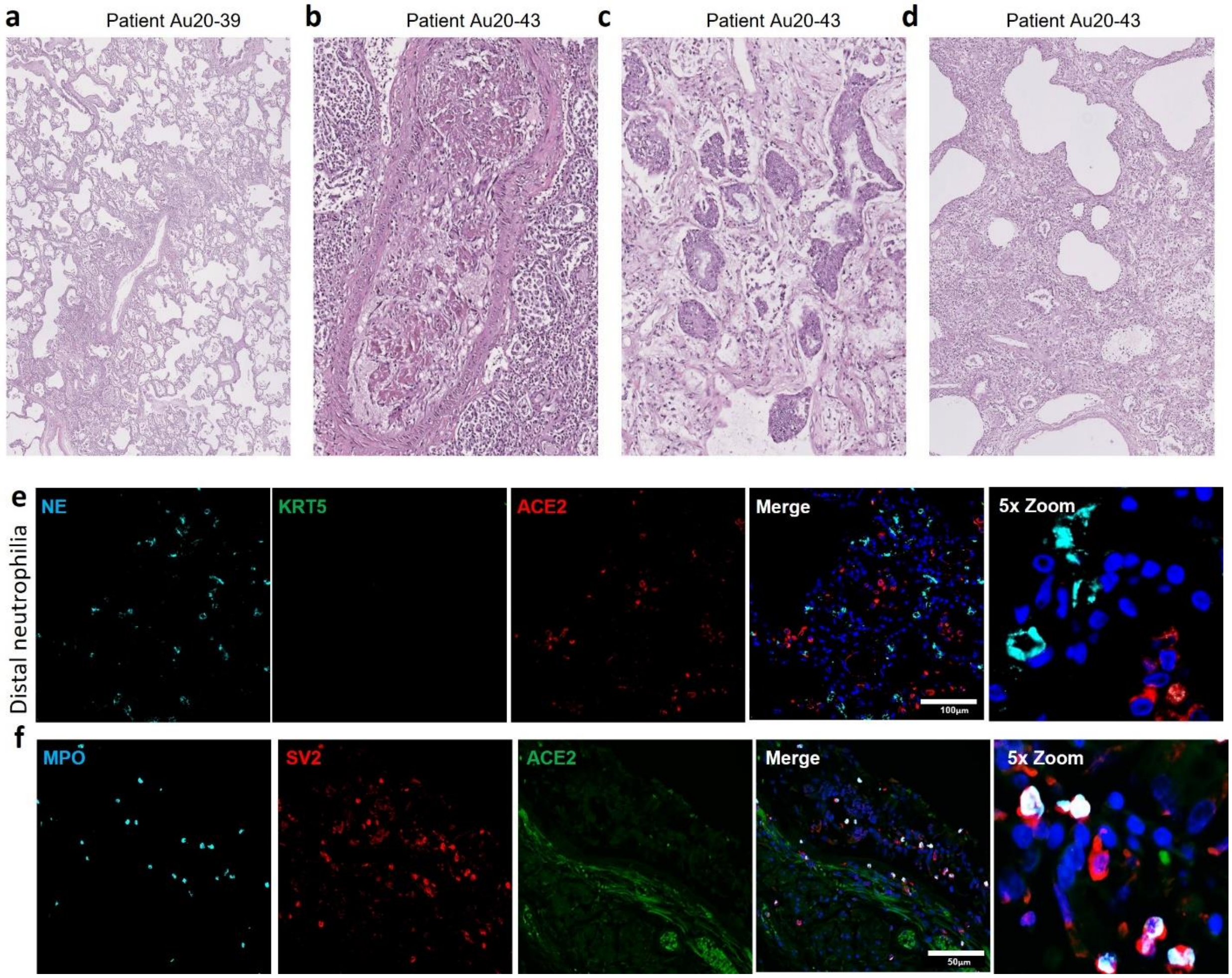

**Supplementary figure S4, related to figure 4: Neutrophil associated tissue pathology in post-mortem COVID19 and underlying respiratory disease in human lung airways.** A-d) Representative H&E staining of postmortem COVID-19 patient tissues (a) patchy organizing pneumonia centered on and artery and adjacent airway (H&E in; original magnification 20x); b) Artery involved by an organizing thrombus. The adjacent alveolar sacs are filled with inflammatory cells including numerous foamy histiocytes (H&E; original magnification 100x). c) Area of peribronchiolar metaplasia with early squamous metaplasia. Peribronchiolar metaplasia is a common finding and may have existed prior to the COVID infection, but the squamous metaplasia appears to be early in phase and is likely a component of the COVID-induced tissue injury (H&E; original magnification 100x). d) Rounded airspaces are within the interstitium and lined by organized inflammatory cells and fibrin, consistent with barotrauma from mechanical ventilation (H&E; original magnification 40x). e) Representative IF image probed for myeloperoxidase (cyan), SARS-CoV-2 (red) and ACE2 (green) All IF images have nuclei counterstained with DAPI (blue) and scale bars represent 50 µm. All images are representative of 3 independent regions per donor at least 2 independent donors.

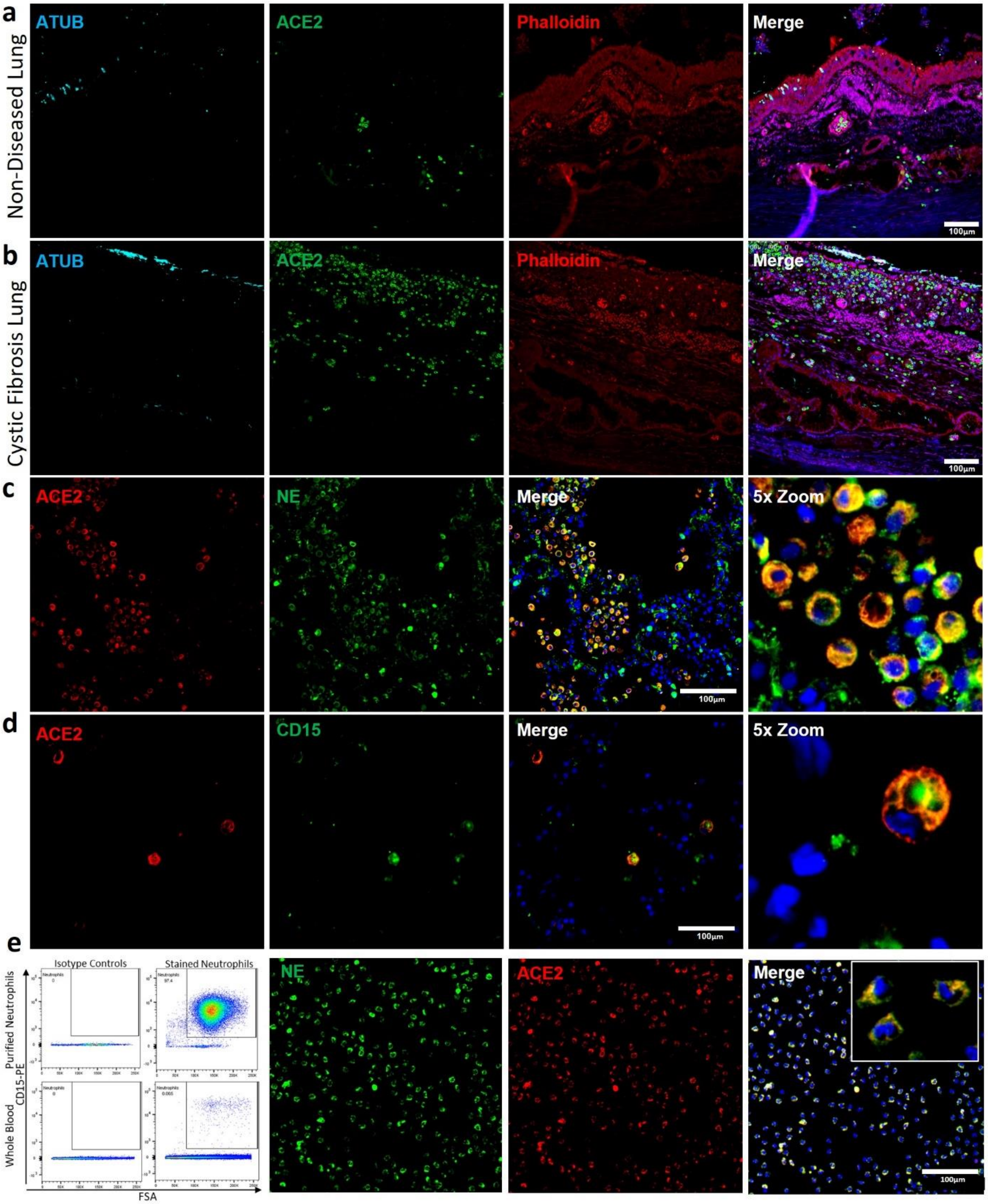

**Supplementary figure S5, related to figure 5. Evidence for ACE2 expression in respiratory infiltrating neutrophils.** A-b) Representative IF images of explanted human lung tissue from no underlying respiratory disease (a) or CF patients (b); for acetylated α-tubulin (cyan), ACE2 (green) and Phalloidin (red). Images highlight; increased ACE2+ Leukocyte infiltrating cystic fibrosis tissue; (c-d) Representative IF images of cystic fibrosis tissue for ACE 2 (red) NE (c) or CD15 (d) (green) demonstrating colocalization of neutrophil markers in cells expressing ACE2. (e) FACS plot and subsequent IF staining of CD15+ Peripheral polymorphonuclear leukocytes isolated from human blood, showing expression of ACE2 in neutrophils. All IF images have nuclei counterstained with DAPI (blue) and scale bars represent 100 μm. All images are representative of 3 independent regions per donor at least 2 independent donors.
